# A chimeric human-mouse lung vascular model using induced pluripotent stem cells reveals insights into the pathogenesis of *BMPR2*-related pulmonary hypertension

**DOI:** 10.64898/2026.04.29.721664

**Authors:** Alexander M. Holtz, Marcus Vorpahl, Mohamed J. Ahmed, Eric D. Austin, Pushpinder S. Bawa, Carlos Villacorta-Martin, Mervin C. Yoder, Darrell N. Kotton

**Affiliations:** Center for Regenerative Medicine, Boston University and Boston Medical Center, Boston, MA 02118 USA; Division of Genetics and Genomics, Boston Children’s Hospital, Boston, MA 02130 USA; Hamdan Bin Mohammed College of Dental Medicine, Mohammed Bin Rashid University of Medicine and Health Sciences, Dubai, UAE; Department of Pediatrics, Vanderbilt University Medical Center, Monroe Carell Jr. Children’s Hospital, Nashville, TN 37204 USA; McGowan Institute for Regenerative Medicine, Department of Surgery, University of Pittsburgh, Pittsburgh, PA 15219 USA; The Pulmonary Center and Department of Medicine, Boston University Chobanian & Avedisian School of Medicine, Boston, MA 02118 USA

## Abstract

Advances in tissue biology have revealed remarkable transcriptomic heterogeneity of endothelial cells between and within organ systems. This necessitates more precise models of organ-specific endothelium to understand the pathogenesis of genetic vascular disorders, such as pulmonary hypertension (PH), where gene-disease associations have implicated endothelial cell dysfunction as a key driver of disease pathogenesis. Towards this end, human induced pluripotent stem cells (hiPSCs) hold immense promise for PH disease modeling where hiPSCs are generated from an affected individual and undergo gene correction to generate syngeneic controls that can be differentiated to endothelial cells (hiEndos), providing a limitless source of material for downstream studies; however, the ability to generate lung-specific hiEndos to model pulmonary vascular disease has been limited. To overcome this challenge, we developed a chimeric human-mouse lung vascular model wherein hiEndos are first patterned via BMP9-induced signaling towards a lung-like molecular phenotype *in vitro* and are then intravenously transplanted into the mouse lung vasculature *in vivo* to generate orthotopic lung-specific endothelium for downstream studies. Transplanted pre-patterned hiEndos form functional connections to the native mouse lung vasculature and upregulate differentiated lung-specific molecular cell subtype profiles that include capillary- and arterial-like cell populations. To apply this approach for disease modeling, we generated new hiPSC lines by reprogramming fibroblasts from individuals of the 2001 landmark cohort of *BMPR2* gene variant-associated PH and developed a novel *in vivo* competitive lung endothelial reconstitution assay to quantify functional and molecular differences between human *BMPR2*-variant vs syngeneic gene-corrected/edited hiEndos. Our approach revealed novel insights into PH disease pathogenesis, not previously evident with prior models, including *BMPR2* variant-induced *in vivo* defects in human lung capillary gene expression, elevated lncRNA *H19* expression, increased AHR signaling, and diminished functional capacity to repopulate the pulmonary vascular endothelium.

## Introduction

Dysfunction of the pulmonary endothelium is a key disease driver in several human acquired and congenital disorders including pulmonary hypertension (PH), which is a devastating disorder with limited treatment options.^1^ Pathogenic variation in genes critical to endothelial cell biology (*BMPR2*, *FOXF1*, etc.) have been associated with both pediatric and adult PH revealing a critical role for the pulmonary endothelium in PH pathogenesis.^2–8^ As the deployment of broad-based sequencing technologies in the clinical space reveals novel gene-disease associations, more precise models of the human lung vasculature are required to clarify the relevance of candidate gene variants, deepen our understanding of disease pathogenesis, and develop novel therapeutic approaches.

Developing more faithful models of the pulmonary vasculature is particularly important in the setting of recent discoveries demonstrating previously unappreciated molecular heterogeneity of endothelial cells between and within organ systems.^9,10^ Single cell RNA sequencing (scRNA-seq) studies in mice and humans show that endothelial cells have distinct transcriptomic signatures depending on their organ of origin, with the capillary endothelium exhibiting the greatest diversity, potentially reflecting their organ-specific function.^9,10^ Within the lung, scRNA-seq has demonstrated two novel capillary cell populations termed CAP1 (also known as general capillary or gCaps) and CAP2 cells (also known as aCaps, Car4 endothelial cells, or aerocytes).^11–14^ CAP2 cells, while less abundant, exhibit large and complex morphology and close apposition to type 1 alveolar epithelial cells (AT1 cells), thus forming the majority of the respiratory surface across which gases exchange between the air and blood.^11,12,14^ By contrast, CAP1 cells demonstrate progenitor capacity as they can proliferate and differentiate into CAP2 cells in response to injury to restore the capillary bed.^11,12^ These novel discoveries necessitate more accurate models of the human lung endothelium to resolve the mechanisms driving pulmonary vascular diseases.

Studies in human PH patients and mouse models have advanced our understanding of PH pathogenesis and the impact of gene variation on vascular biology. Genome wide association studies followed by refined mutation mapping identified variation in the *BMPR2* gene as the first gene-disease association in PH in familial and sporadic PH.^2,4,5,7,15,16^ Subsequently, mouse models of *BMPR2* haploinsufficiency did not spontaneously develop PH, but did show an exaggerated disease response in certain PH models, while conditional knockout of *BMPR2* in endothelial cells was sufficient to generate PH pathology.^17–21^ This suggests that mice may be more resilient to *BMPR2* haploinsuffiency and that advanced human disease models are needed to unravel *BMPR2*-related PH pathogenesis and to clarify variant pathogenicity.

Human induced pluripotent stem cells (hiPSCs) have served as a powerful tool for understanding the impact of pathogenic gene variation on PH pathogenesis in a human cell context.^22–25^ hiPSCs derived from individuals with a genetic disorder, such as hereditary PH, possess the key pathogenic variant in addition to the individual’s entire genetic background. CAS9-mediated gene correction can be used to generate syngeneic control cells and the relevant candidate cell types contributing to disease pathogenesis can be derived through directed differentiation to provide a limitless source of cells for downstream studies. This is particularly relevant for human lung disease where primary tissue from diseased individuals is extremely limited, particularly at early stages of disease, and cells within these tissues are not amenable to manipulation for functional studies. Prior studies utilizing hiPSC-derived endothelial cells (hiEndos) from patients with *BMPR2*-related PH, reported in vitro defects in endothelial adhesion, migration, angiogenesis, and apoptosis in 2D culture systems;^23,24^ however, it remains unclear whether the endothelial-like cells generated express lung-specific molecular and functional programs as the derived cells have not experienced the typical intercellular interactions, inter-organ crosstalk, and biophysical forces encountered by native lung endothelium. Since *BMPR2*-deficient endothelial cells cause clinical disease most conspicuously in the pulmonary vasculature, pattering hiEndos into lung-specific endothelial lineages *in vitro* is likely to be important for disease modeling, and developing novel systems that allow their study in a native tissue context *in vivo* may address these limitations and provide deeper insights into PH pathogenesis.^9,10^

To overcome these challenges, we present here a novel *in vivo* chimeric human-mouse lung vascular model to interrogate the mechanisms of genetic pulmonary vascular disease. By first inducing hiEndos *in vitro* to express lung-specific programs we successfully engraft either normal or disease-specific human cells into the mouse lung vasculature. Our novel directed differentiation and expansion protocol rapidly generates billions of hiEndos *in vitro* that can undergo freeze-thaw and extensive passaging in culture, while maintaining proliferative capacity and endothelial cell fate. To quantitatively assess the fitness of various hiEndos to engraft in the mouse lung vasculature *in vivo*, we develop a competitive lung endothelial reconstitution assay and observe that pre-treatment of hiEndos with BMP9 prior to transplantation shifts cells towards a lung-specific molecular profile and promotes their lung engraftment. Engrafted-hiEndos form functional connections to the native mouse lung vasculature and further augment lung-specific transcriptional profiles differentiating into both capillary- and arterial-like populations. Finally for *in vitro* and *in vivo* disease modeling we generate hiPSC lines from archival samples of the landmark *BMPR2*-related PH cohort^7^, edit the *BMPR2* variant locus to produce paired syngeneic corrected hiPSC clones, and then define the molecular phenotypes of their hiEndo progeny, both *in vitro* and after competitive reconstitution of the mouse lung vasculature *in vivo*. We find competing hiEndos carrying *BMPR2* variants exhibit significantly diminished *in vivo* competitive repopulation capacity compared to their gene-corrected counterparts, and transcriptomic profiling of the engrafted pairs reveals novel insights into disease pathogenesis including deficiencies in CAP2 gene expression, aberrant lncRNA *H19* expression, and elevated AHR signaling.

## Results

### Stable expansion of hiEndos in culture through combined inhibition of Notch and TGF-beta signaling

To generate the large numbers of hiPSC-derived endothelial cells (hiEndos) required for both disease modeling and transplantation, we first sought to develop a differentiation protocol in defined conditions that would allow extensive expansion of human cells through both freeze-thaw as well as serial passaging cycles, while maintaining endothelial cell phenotype. We modified previously published methods^26,27^ to differentiate hiPSCs generated from multiple normal donors (BU1 line^28^, Figure 1; and BU3 line^29^, Figure S1) through the early stages of gastrulation (primitive streak followed by lateral plate mesoderm), an approach known as “directed differentiation”. As expected based on prior publications, we observed that cells treated for 24 hours with a combination of BMP4, FGF2, and the GSK3-beta inhibitor, CHIR99021, generated T-brachyury expressing primitive streak-like cells (Figure 1A-C, S1A). Subsequent treatment for 24 hours with a combination of BMP4 and FGF2 induced differentiation towards lateral plate mesoderm, as evidenced by induction of *KDR* and the lateral plate mesoderm marker, *FOXF1,* without expression of the endoderm marker *FOXA2* (Figure 1 A-D, S1A). While we observed an initial induction of intermediate and paraxial mesoderm markers *PAX2* and *TBX6,* respectively, expression of these markers decreased throughout the differentiation while high levels of *FOXF1* expression was maintained (Figure 1 C and D, S1B). Next, we induced endothelial lineage specification of mesoderm through combined treatment with high-dose VEGF (200ng/mL) and the Notch inhibitor DAPT.^30^ We observed expression of endothelial cell marker transcripts, *PECAM1* and *CDH5,* starting 24 hours after endothelial cell induction, with increasing expression over time (Figure 1E, S1C). To monitor expression efficiency of these markers at the protein level, we performed flow cytometry on day 5 of differentiation, finding an average of 21.9% (range 10-60%) of cells co-expressed CD31/CD144 (Figure S1D). We purified day 5 hiEndos for characterization and expansion using magnetic activated cell separation (MACS) with CD31 and CD144 microbeads (Figure S1E). The purified cells exhibited co-expression of CD31 and CD144 protein by flow cytometry and immunofluorescence microscopy, a cobblestone morphology on 2D Matrigel, capillary network formation when plated on 3D Matrigel, and uptake of fluorescently labeled acetylated low-density lipoprotein (AcLDL, Figures 1B, F-J, S1F and G), features commonly reported as associated with cultured endothelial cell phenotypes.^26,27,30–32^ While we successfully and rapidly generated hiEndos from multiple hiPSC lines, we found purified hiEndos to be unstable after serial passaging in these culture conditions, as evidenced by diminishing frequencies of CD31/CD144 co-expression, emergence of cells with a fibroblast morphology coincident with loss of endothelial cobblestone morphology, and increasing expression of fibroblast markers *PDGFRA* and *COL1A1*, but not smooth muscle markers *MYH11* or *ACTA2* (Figure S1H-J).

**Figure 1.**
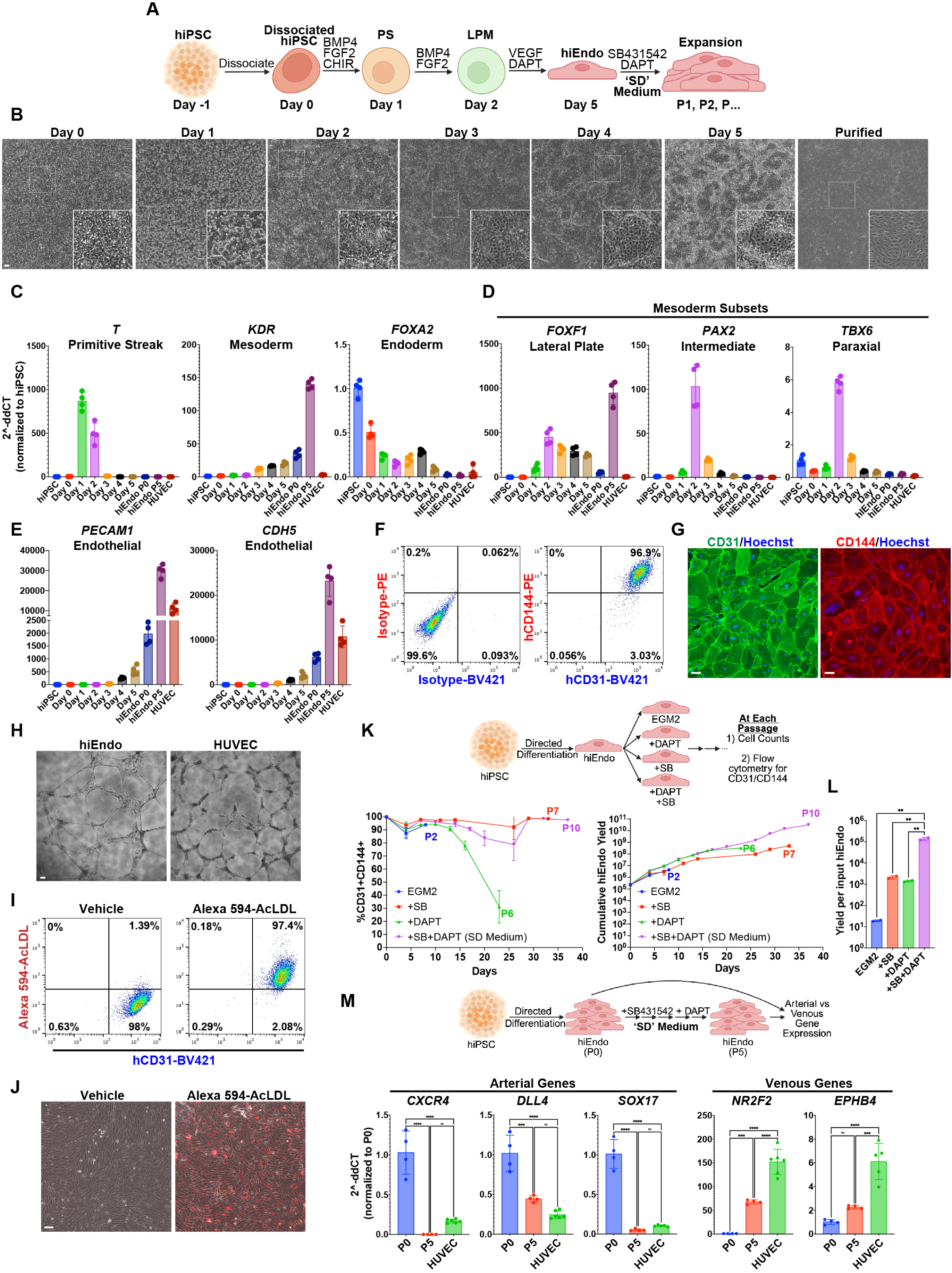
Directed differentiation strategy to generate hiEndos that are expanded using combined inhibition of Notch and TGF-beta signaling. (A) Schematic of directed differentiation protocol to generate hiEndos. (B) Phase contrast images during the indicated day of differentiation. White dashed square represents region magnified in insets. Scale bar: 50um. (C-E) RT-qPCR for the indicated primers using RNA isolated from cells on each day of differentiation, purified hiEndos at P0 and P5, and HUVECs. N=4 experimental replicates from independent wells of the same differentiation, representative results from two independent differentiations. (F) Flow cytometry of hiEndos using antibodies against the human endothelial lineage markers CD31 and CD144 (right) and isotype controls (left), N=4 biological replicates from distinct differentiations. (G) Immunostaining of hiEndos for human CD31 (green, left) and CD144 (red, right). Hoechst stains nuclei (blue). Scale bar 10um. N=3 biological replicates. (H) Capillary tube formation assay on 3D Matrigel. Scale bar 50um. N=4 biological replicates. (I-J) Human CD31 and acetylated LDL uptake in hiEndos measured by flow cytomtetry (I) or fluorescence microscopy (J). Scale bar 50um. N=8 biological replicates. (K) Schematic of experiment to test the impact of Notch (DAPT) and TGF-beta (SB431542) inhibition on hiEndo expansion (top). Quantification of CD31 and CD144 double-positive cells by flow cytometry (left) and cumulative hiEndo yield (right) at each passage grown in the indicated media conditions. N=2 experimental replicates from independent wells of the same differentiation, representative data shown from 3 independent differentiations. (L) Quantification of total hiEndo yield per input hiEndo in each media. ** p<0.01 using one-way Anova and Tukey’s multiple comparison test. (M) RT-qPCR of arterial and venous markers after serial passaging in SD medium. N=4 experimental replicates from independent wells of the same differentiation, representative results from two independent differentiations. ** p<0.01, *** p<0.001, **** p<0.0001, one-way Anova and Tukey’s multiple comparison test. Abbreviations: hiPSC (human induced pluripotent stem cell), CHIR (CHIR99021), P (passage), HUVEC (human umbilical vein endothelial cells), AcLDL (acetylated low-density lipoprotein), SB (SB431542).

Instability of endothelial phenotypes in culture has been extensively reported and significantly limits applications that require large cell numbers, such as disease modeling and transplantation.^32–35^ To address this, we next aimed to develop methods for stable *in vitro* expansion of our purified hiEndos while maintaining their endothelial phenotype. We reasoned that TGF-beta inhibition with SB431542 would maintain endothelial phenotype in culture, similar to previous approaches.^33^ Additionally, we reasoned that inhibition of Notch signaling with DAPT would maintain hiEndo proliferative potential given the role of Notch signaling in endothelial cell maturation.^36^ To determine if we could enhance hiEndo cumulative yield over time, we measured hiEndo efficiency and yield at each passage in cells grown with vehicle, SB431542, DAPT, or combined SB431542 and DAPT (Figure 1K). Treatment with SB431542 alone maintained hiEndo cell fate longer than untreated cells, but we observed growth arrest after several passages (Figure 1K). Notch inhibition alone increased cell yield compared to untreated cells; however, we still observed loss of endothelial cell fate after serial passaging (Figure 1K). In marked contrast, combined treatment with SB431542 and DAPT preserved hiEndo cell fate and proliferative potential across serial passages, yielding billions of CD31/CD144 double positive hiEndos after ∼1 month in culture (Figure 1K). This combined inhibition strategy increased the yield of hiEndos per input cell by two orders of magnitude compared to SB431542 or DAPT treated cells (Figure 1L). We also found that hiEndos grown in SB431542 and DAPT medium (herafter “SD”) could maintain their cell fate, proliferative potential, and passageability after archival freezing and thawing (data not shown), a key milestone needed to maintain stable cell archives for downstream *in vitro* or *in vivo* large-scale applications.

To determine the arterial vs venous nature of hiEndos before vs after culturing in SD medium, we compared the expression of lineage markers in newly purified day 5 hiEndos compared to those that have been transitioned to and passaged in SD medium. We reasoned that although high dose VEGFA used in our endothelial specification medium has been shown to promote arterial identity,^32^ ongoing Notch inhibition with DAPT is likely to promote venous identity over time, given the known role of Notch signaling to generate arterial cell fates.^32,36^ Consistently, we observed initial expression of the arterial markers *CXCR4*, *DLL4*, and *SOX17* in P0 hiEndos, but their expression decreased over serial passages, while expression of venous markers *NR2F2* and *EPHB4* were initially low and increased after growth in SD medium (Figure 1M). These data suggest that day 5 hiEndos exhibit cell fate plasticity to adopt arterial vs venous identities depending on the subsequent culture conditions, and serial passaging in SD medium increases venous gene expression at the expense of arterial gene marker expression.

### BMP9 and VEGF modulates lung-specific gene expression in hiEndos

We next explored the competency of hiEndos to adopt a lung endothelial-specific gene expression profile *in vitro*. Single-cell RNA sequencing (scRNA-seq) studies have identified *TMEM100* as a unique marker of lung endothelium in both adult mice and humans (Figure 2A).^9,10,37^ *TMEM100* is a down-stream target of BMP9 signaling in endothelial cells so we first explored whether *TMEM100* expression in hiEndos could be induced by BMP9 treatment.^33^ Indeed, we found that BMP9 treatment increased *TMEM100* expression in hiEndos in a dose-dependent manner and up-regulated the canonical BMP target genes *ID1* and *ID2* (Figures 2B and S2A). To determine whether BMP9 impacts expression of other genes related to lung endothelial cell identity, we measured expression of CAP1 and CAP2 gene markers in hiEndos in response to BMP9 treatment (Figure 2B-G and S2). We found that BMP9 decreased expression of 5 out of 6 CAP2 markers assayed (including *EDNRB*, *HPGD, KITLG*, *CA4*, and *APLN*; Figure 2B-G and S2B-D), with mixed effects on CAP1 gene expression (increasing 3 out of 5 markers assayed: *GPIHBP1, FCN3, and CD36;* Figure 2B-G and S2B-E). This suggests that BMP9 treatment, while promoting expression of the lung specific marker, *TMEM100*, may be inhibitory towards CAP2 identity.

**Figure 2.**
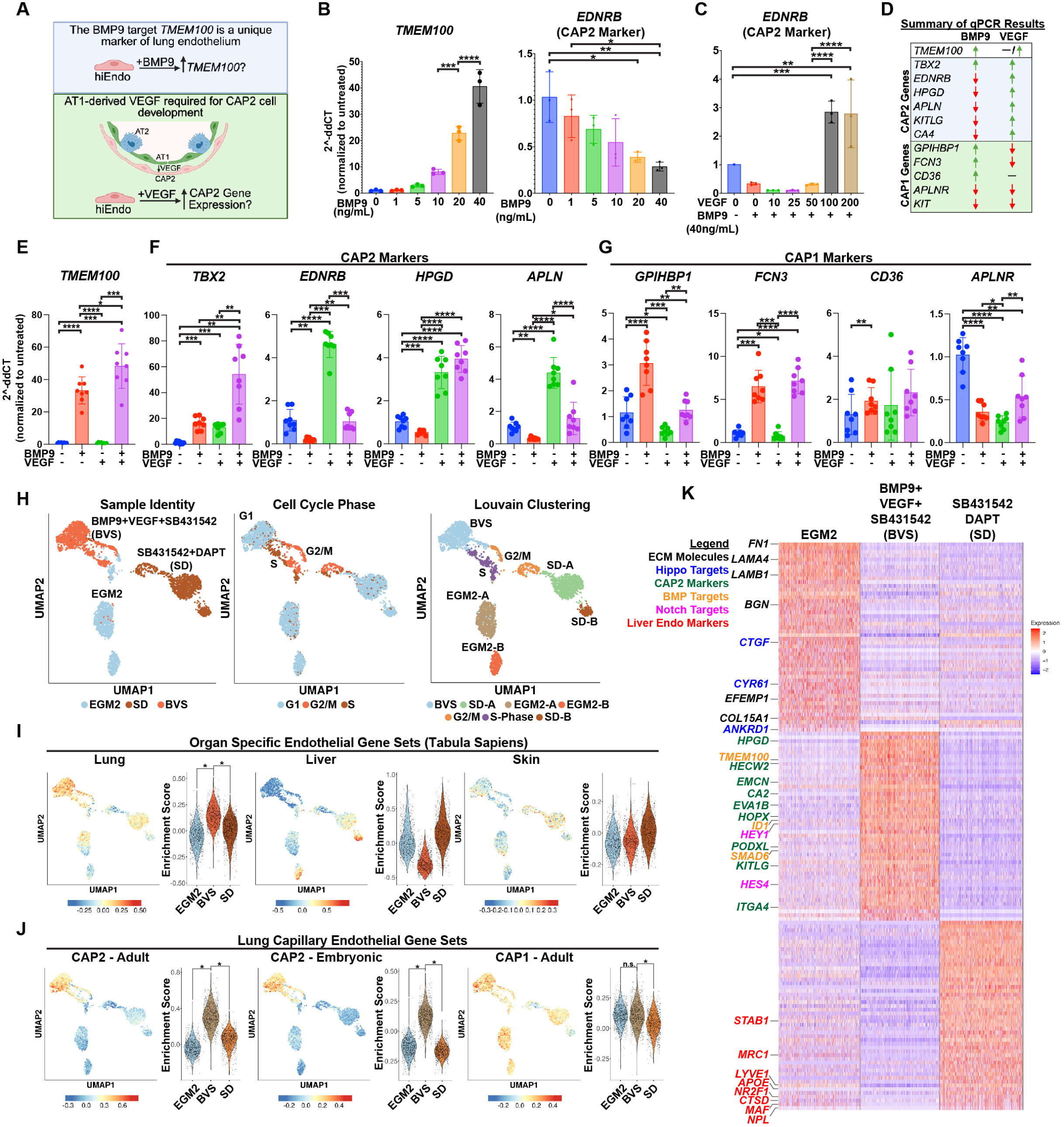
Modulation of lung capillary endothelial gene expression in hiEndos by BMP9 and VEGF. (A) Schematic of rationale for testing BMP9 and VEGF treatment of hiEndos to modulate lung endothelial gene expression. (B) RT-qPCR analysis of *TMEM100* and *EDNRB* expression in hiEndos treated for 72 hours with varying concentrations of BMP9. (C) As in (B) except with varying concentrations of VEGFA in combination with BMP9. N=3 experimental replicates from independent wells of the same differentiation, representative data shown from three distinct differentiations. * p<0.05, ** p<0.01, *** p<0.001, **** p<0.0001, one-way Anova and Tukey’s multiple comparisons. (D) Summary of RT-qPCR results from (E-G) and Figure S2. (E-G) Fold change in expression of indicated genes (RT-qPCR) after treatment with the indicated growth factors for 7-days. N=8 biological replicates per condition from distinct differentiations. * p<0.05, ** p<0.01, *** p<0.001, **** p<0.0001, one-way Anova with Tukey multiple comparisons. (H) UMAP representation of scRNA-seq profiles of hiEndos grown in the indicated media conditions for 7-days showing sample identify (left), cell cycle phase (middle), and Louvain clustering (right). (I-J) Enrichment scores represented on UMAP and violin plots using gene sets for organotypic endothelial identity (I)^9^ or CAP1/2 cells (J).^13,53^ * p<1e-50. (K) Heat map shows the top 50 differentially expressed genes in each culture media. Select genes are highlighted and their category indicated by color. Abbreviations: BVS (BMP9+VEGFA+SB431542), SD (SB431542+DAPT)

Although little is known regarding CAP2 specification in humans, recent evidence implicates juxtacrine VEGFA signaling from AT1 cells as an inducer of CAP2 cell fate in mice.^14^ Therefore, we investigated whether VEGFA treatment may positively regulate CAP2 gene expression in hiEndos in the context of BMP9 treatment (Figure 2A). To test this, we measured CAP2 gene expression in hiEndos treated with BMP9 together with increasing concentrations of VEGFA. We observed a VEGFA dose-dependent increase in the expression of the CAP2 gene markers *EDNRB*, *TBX2*, *CA4*, and *APLN* compared to cells treated with BMP9 alone (Figures 2C and S2C), consistent with a role for VEGFA in promoting CAP2 cell fate. To directly compare the impact of VEGFA and BMP9 treatment on lung endothelial gene expression in hiEndos when added to our base medium of EGM2+SB431542, we compared CAP1 and CAP2 gene expression in cells treated with vehicle, BMP9 (40ng/mL), VEGFA (200ng/mL), or combined treatment with BMP9 and VEGFA. The results of these studies are summarized in Figure 2D and detailed in Figures 2 and S2. First, while VEGFA alone had no effect on *TMEM100* expression, we observed a significant increase in *TMEM100* expression with combined BMP9 and VEGFA treatment compared to BMP9 treatment alone (Figure 2E). We next found that both BMP9 alone and VEGFA alone up-regulated the CAP2 marker *TBX2* and that combined treatment led to a synergistic increase in expression (Figure 2F). With respect to other CAP2 markers, we confirmed that BMP9 treatment reduced the expression of *EDNRB*, *HPGD*, *APLN*, *KITLG*, and *CA4* while VEGFA increased expression of all CAP2 markers tested (Figures 2F and S2D). Note, we observed variability in the extent of *EDNRB* expression above baseline in response to combined BMP9 and VEGFA treatment (Figure 2C and F). With respect to CAP1 markers, we found that BMP9 treatment induced expression of CAP1 markers, *GPIHBP1*, *FCN3*, and *CD36* while inhibiting expression of *APLNR* and *KIT* (Figure 2G and S2E). By contrast, VEGFA treatment reduced all CAP1 markers except for *CD36* (Figure 2G and S2E). Overall, these results demonstrate that BMP9 and VEGFA play overlapping and opposing roles to modulate CAP1 and CAP2 gene expression with VEGFA promoting CAP2 and inhibiting CAP1 gene markers.

Having established that BMP9+VEGFA added to our SB431542+EGM2 medium (hereafter denoted “BVS”) synergistically augmented expression of the endothelial specific marker, *TMEM100*, we next investigated the molecular profiles and heterogeneity of cells treated with BVS compared to base (EGM2) or SD media by scRNA-seq. We found that hiEndo transcriptomes clustered largely by culture medium treatment (Sample Identity) and cell cycle phase (Louvain; Figure 2H) with less heterogeneity in clustering for BVS-treated hiEndos in the G1 cell cycle phase, compared to cells in other media. To determine whether BVS medium induced a lung endothelial transcriptional program, we quantified expression levels of gene modules that define organotypic endothelial identity derived from the published atlas, Tabula sapiens.^9^ This analysis demonstrated enrichment of a lung endothelial gene set in BVS-treated cells compared to EGM2 and SD media, but no enrichment in liver or skin endothelial gene sets (Figure 2I, see Table S1 for gene sets). This demonstrated the specificity of BVS medium to augment lung-specific gene expression (Figure 2I). In contrast, EGM2-B and SD-B cell clusters were enriched for a liver-specific endothelial gene set (Figure 2I), further indicating that modulating signaling pathways in hiEndo outgrowth media can regulate the tissue patterning logic of endothelial fated cells.

Beyond promoting lung endothelial patterning, based on our RT-qPCR studies (Figures 2 and S2) we reasoned that BVS medium would induce a CAP2 gene expression profile. Therefore, we characterized CAP1 and CAP2 gene set expression in each media, finding near uniform enrichment of adult and embryonic CAP2 gene sets in BVS-treated cells compared to other conditions (Figure 2J), without differential enrichment of a CAP1 gene set (Figure 2J). These data collectively demonstrate that BVS medium induces both lung endothelial patterning and CAP2 gene expression in hiEndos and that the majority of hiEndos possess competence to respond to this treatment with minimal heterogeneity.

To gain a deeper understanding of the cellular responses to the different media conditions, we next performed differential gene expression analysis. We found enrichment of the BMP-target genes *TMEM100*, *ID1*, and *SMAD6* in addition to CAP2-specific genes in the top 50 differentially expressed genes (DEGs) in BVS-treated hiEndos (Figure 2K, Table S2).^13,34^ We also identified up-regulation of the Notch target genes *HEY1* and *HES4* in BVS-treated hiEndos consistent with prior studies demonstrating cross-talk between BMP9 and Notch signaling.^38,39^ Cells grown in EGM2 base medium alone exhibited enrichment of extracellular matrix molecules (*FN1*, *LAMA4*, *LAMB1, COL15A1*) and Hippo pathway targets (*CTGF*, *CYR61*, and *ANKRD1*) that may reflect cooperativity between TGF-beta and Hippo signaling to regulate endothelial-to-mesenchymal transition (Figure 2K, Table S2).^40,41^ This analysis also revealed liver endothelial-specific genes in the top DEGs in hiEndos treated with SD medium consistent with our gene set analysis (Figure 2K, Table S2). We next performed gene set enrichment analysis (GSEA) to gain further insight into candidate pathways altered in the different media conditions (Figure S2F). Consistent with the above analysis, we found that cells in EGM2 medium exhibited an increased response to TGF-beta signaling (Figure S2F, Table S3). BVS-treated hiEndos showed an increase in genes related to oxidative phosphorylation and cells grown in SD medium showed an increase in genes related to cytoplasmic translation (Figures S2F, Tables S4 and S5).

### BMP9 promotes engraftment of hiEndos in the injured mouse lung

Given the competence of hiEndos to respond to key lung endothelial signaling molecules *in vitro*, we sought to test whether this patterning in culture would impact the molecular or functional phenotypes of the cells *in vivo* following orthotopic transplantation. Given the molecular heterogeneity observed between organ-specific endothelia, we hypothesized that ‘patterning’ hiEndos towards a lung-specific molecular profile through treatment with BMP9 *in vitro* would promote lung engraftment after intravenous delivery into NSG mouse recipients. To test this hypothesis, we first developed a competitive lung endothelial reconstitution assay (Figure 3A) to obtain a quantitative *in vivo* readout of the engraftment fitness of hiEndos undergoing distinct pre-treatment strategies, similar to classical experiments assaying the fitness of candidate hematopoietic stem cell populations to reconstitute the bone marrow, reported previously as “competitive blood repopulation”.^42,43^ We lentivirally transduced hiEndos *in vitro* to constitutively express either EGFP or dsRED to track cells following transplantation.^44,45^ Our stepwise strategy for developing this assay was to compete these differentially labelled hiEndo populations after exposing them to distinct pre-treatment strategies, followed by mixing the populations 1:1 for retro-orbital injections into NSG mice that were pre-exposed to 72 hours of hyperoxia to damage the native lung microvasculature, a conditioning strategy that has been previously used to accommodate primary lung endothelial cell transplantation.^46,47^

**Figure 3.**
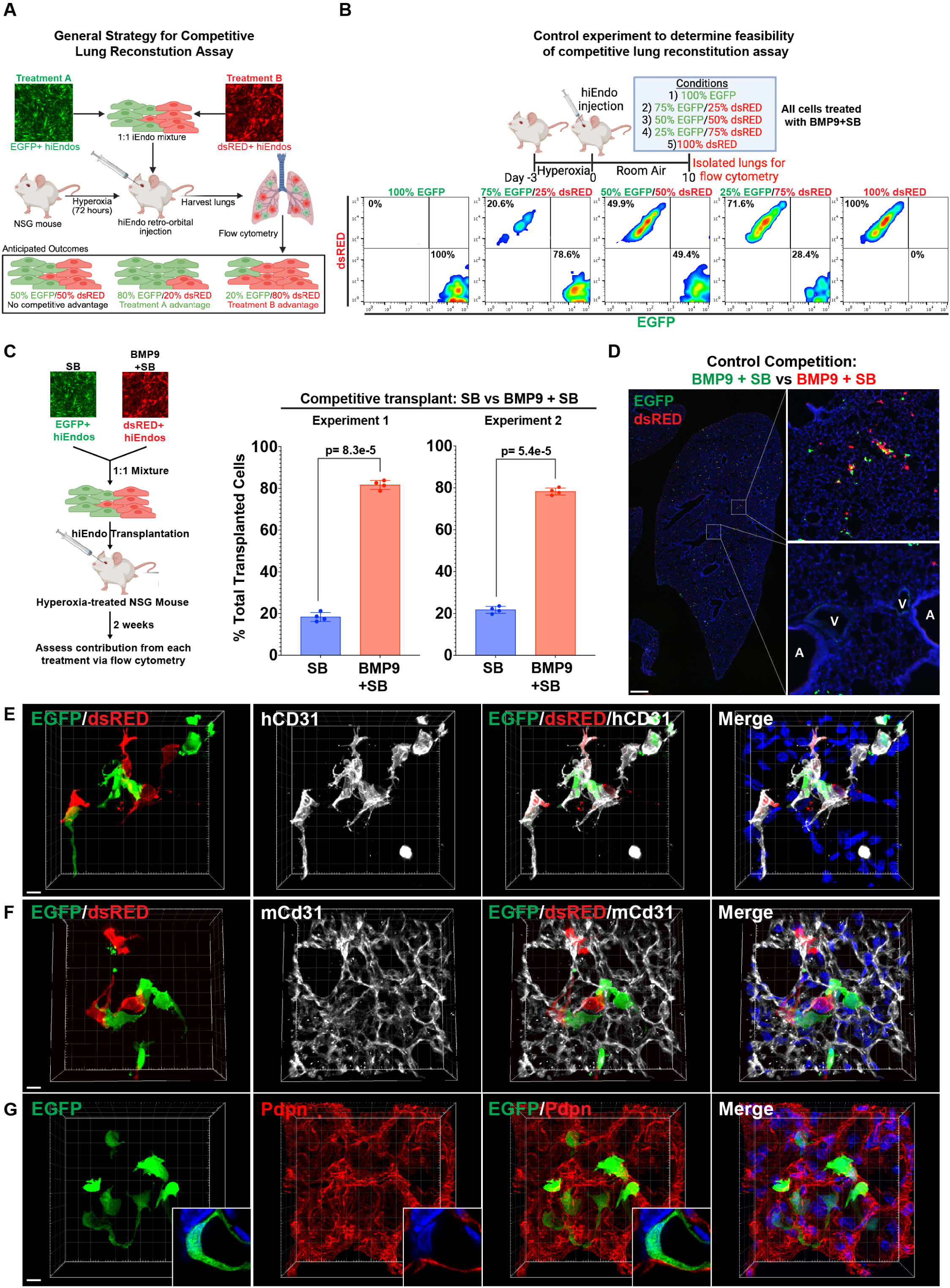
BMP9 promotes hiEndo engraftment in continuity with the mouse lung vasculature. (A) Schematic of the general strategy for the competitive lung reconstitution assay. (B) Cartoon depiction of experiment to test feasibility of competitive lung reconstitution assay by transplanting defined ratios of EGFP+ and dsRED+ cells that were all pre-treated with BMP9+SB431542 (top). Flow cytometry gated on human CD31+ post-transplant hiEndos 10-days following injection and analyzed for dsRED and EGFP expression (bottom). N=1 animal per condition. (C) Quantitation of the contribution of SB- vs BMP9+SB-treated hiEndos in the mouse lung 2 weeks following transplantation as determined by flow cytometry across 2 distinct experiments. N=4 animals per experiment, p-value determined by Student’s paired two-tailed t-test. (D) Left lung lobe immunofluorescence microscopy to identify EGFP+ and dsRED+ hiEndos 2-weeks following transplantation of BMP9 pretreated hiEndos. hiEndos are largely present in the alveolar space (top inset) as opposed to larger vessels (bottom inset). White dotted lines indicate region of inset. A airway, V vessel. N=3 animals. (E-G) Immunofluorescence confocal microscopy of recipient mouse lung tissue sections 2 weeks following transplantation of mixtures of EGFP+ and dsRED+ hiEndos. 3D reconstruction of Z-stacks shows each indicated antibody-stained fluorochrome together with immunostaining for human CD31 (hCD31, E, white), mouse Cd31 (mCd31, F, white), or podoplanin (Pdpn, G, red). Insets (G) show close apposition of Pdpn and EGFP+ hiEndo. Hoechst stains nuclei (blue). Scale bars (E-G) 10um. N=4 animals.

Since there have been very few prior attempts to functionally engraft human cells of any kind in the pulmonary vasculature^48,49^ we first sought to qualitatively determine whether pulmonary engraftment of hiEndos is feasible. We intravenously injected hiEndos pre-treated with either SB431542 alone, or combined BMP9+SB431542 into separate cohorts of immunodeficient NSG mice. While few to no labeled cells were observed in uninjured recipients, labeled cells were easily visualized in the lung parenchyma of recipients pre-conditioned with 72 hours of hyperoxia exposure (Figure S3A). We also observed a qualitative increase in labeled cell presence in recipients of hiEndos that had been pre-treated with BMP9 compared to SB431542 alone (Figure S3A). Having determined that hyperoxia pre-conditioning of NSG recipients and BMP9+SB431542 treatment of hiEndos prior to intravenous delivery provides a scoreable signal for orthotopically transplanted human cells, next we sought to adapt this approach to develop quantitative competitive lung endothelial reconstitution. We competed equally-treated hiEndos, labeled with distinct fluorochromes (EGFP vs dsRed) which were then mixed in varying ratios at the time of IV transplantation (mixed 3:1; 1:1: 1:3, or single-colored controls). We found that the ratios of EGFP:dsRED expressing cells (all pre-treated with BMP9+SB431542) at the time of transplantation were maintained in recipient lungs 10 days post-transplantation (scored by flow cytometry of dissociated lungs; Figure 3B). These data collectively demonstrate the feasibility of the competitive lung endothelial reconstitution assay to obtain a quantitative readout of cellular fitness for transplantation.

We next aimed to apply this new quantitative *in vivo* assay to competitively measure the engraftment fitness of cells treated with varying factors. We first repeated testing of hiEndos pre-treated with BMP9+SB431542 compared to those treated with SB431542 alone, but this time using our competitive lung endothelial reconstitution assay. We found the addition of BMP9 quantitatively increased transplantation efficiency ∼5-fold (Figure 3C).^39,40^ Furthermore, pre-treatment with VEGFA, BMP4, CHIR, or FGF2 in addition to BMP9+SB431542 failed to increase transplantation fitness compared to cells treated with BMP9+SB431542 alone (Figure S3B). To determine the overall contribution of transplanted hiEndos to the lung vasculature, we assayed the relative abundance of hiEndos to mouse endothelial cells in dissociated lungs 2-weeks post transplantation by flow cytometry. We found an average of ∼1.6% of the lung endothelium was composed of human hiEndos at this time point (Figure S3C). Overall, these data validate the competitive lung endothelial reconstitution assay and demonstrate that BMP9 pre-treatment increases the lung transplantation fitness of hiEndos.

Next we assessed whether the transplanted hiEndos structurally and functionally contributed to the pulmonary vasculature *in vivo,* key measures needed to determine whether these exogenous cells had truly engrafted. We first investigated the spatial localization of donor hiEndos within the recipient mouse lung. Whole lobe imaging of tissue sections from mice injected with BMP9+SB431542 pre-treated dsRED+ and EGFP+ hiEndos demonstrated a broad distribution of transplanted cells throughout all lung lobes with predominant localization in alveolar tissue and not in larger vessels (Figure 3D). Confocal microscopy with 3D reconstruction revealed that transplanted hiEndos adopted a complex morphology and maintained expression of human CD31, demonstrating maintenance of cell fate following transplantation (Figure 3E). Transplanted hiEndos were found in alveolar septae in close proximity to endogenous mouse capillary endothelial cells and in close apposition with PDPN-expressing AT1 cells, similar to the localization of endogenous CAP2 cells (Figure 3F and G, insets). These data demonstrate that transplanted hiEndos localize to the alveolar space, structurally resemble microvascular endothelial cells, and maintain their endothelial cell fate *in vivo*.

To determine whether transplanted hiEndos were functionally engrafted in continuity with the existing mouse lung microvasculature, we intravenously administered fluorophore-conjugated Ulex Europaeus Agglutinin I (649-UEAI), a lectin that binds to human endothelial glycocalyx, or vehicle (PBS) into hiEndo-transplanted mouse recipients. We reasoned that this intravascular dye would label transplanted cells only if they experienced blood flow via engraftment to the native lung microvasculature (Figure S3D). With this approach, we found that the majority of transplanted hiEndos were labeled with intravenously administered 649-UEAI as determined by flow cytometry (Figure S3E and F) and confocal microscopy (Figure S3G-I), demonstrating functional engraftment in continuity with the native mouse lung microvasculature.

### Lung-engrafted hiEndos exhibit lung organotypic gene expression in vivo

We next investigated whether hiEndos engrafted in the mouse lung exhibit lung-specific gene expression by scRNA-seq. hiEndos were isolated from mouse lungs 5-weeks post-transplantation via fluorescent activated cell sorting (FACS) for EGFP and hCD31 expression (Figure 4A). More than 90% of EGFP+ cells were hCD31+ suggesting that transplanted hiEndos maintained their cell fate 5-weeks post-transplantation (Figure S4A). Consistent with these results, post-transplant hiEndos exhibited expression of key endothelial lineage marker transcripts including *PECAM1*, *CDH5*, *and KDR* (Figure S4B). Despite *TMEM100* expression being selectively enriched in a subset of hiEndos, the majority of transplanted hiEndos were enriched for a lung-specific endothelial gene set in contrast to a liver endothelial gene set suggesting their lung identity (Figure S4C and Figure 4B). Louvain clustering identified 5 distinct cell clusters (Figure 4C; Supplemental Table 1), and we annotated two of these as lung capillary endothelial cell 1 and 2 (iLCapEC1/2) based on differential gene expression analysis that revealed: 1) expression of CAP1 and CAP2 markers in the top 20 DEGs between cell clusters (Figure 4D, Table S6); 2) enrichment of CAP1 and CAP2 gene sets (Figure 4E); and 3) expression of select markers of CAP1 and CAP2 populations (Figures 4F). While both capillary populations expressed mixed CAP1 and CAP2 marker transcripts, we noted a dichotomous expression of *APLNR* and *APLN* in iLCapEC1 and iLCapEC2 populations, respectively, reminiscent of CAP1 and CAP2 endothelial cells *in vivo* (Figure 4D-F, Table S6).^11,12,14^ We annotated a third cluster as induced lung arterial endothelial cell (iLArtEC), based on differential gene expression analysis showing arterial-specific gene expression in the top 20 DEGs (Figure 4D, Table S6), enrichment of an arterial gene set in this population (Figure 4E), and expression of select arterial (Figure 4F) but not venous genes (Figures 4E,F).^11,12,14^ We annotated a fourth cell cluster as VEGFA+ iLEC based on similar lung capillary gene expression with high *VEGFA* expression (Figure 4C-F, Figure S4D). Lastly, a small cluster of proliferative hiEndos were annotated as proliferating iLEC based on expression of proliferation markers (Figure 4C, Table S6). RNA *in situ* hybridization validated the presence of human *APLNR*+ iLCapEC1, *APLN*+ iLCapEC2, *VEGFA*+ iLEC, and *GJA4+* iLArtEC cells which co-expressed human *PECAM1* and localized to alveolar regions of the distal lung (Figures 4G and H). Overall, these data reveal engrafted hiEndos express heterogeneous transcriptional programs reminiscent of several lung-related vascular subtypes, and taken together with their structural and functional contributions to the vasculature they represent a chimeric human-mouse lung *in vivo* vascular model.

**Figure 4.**
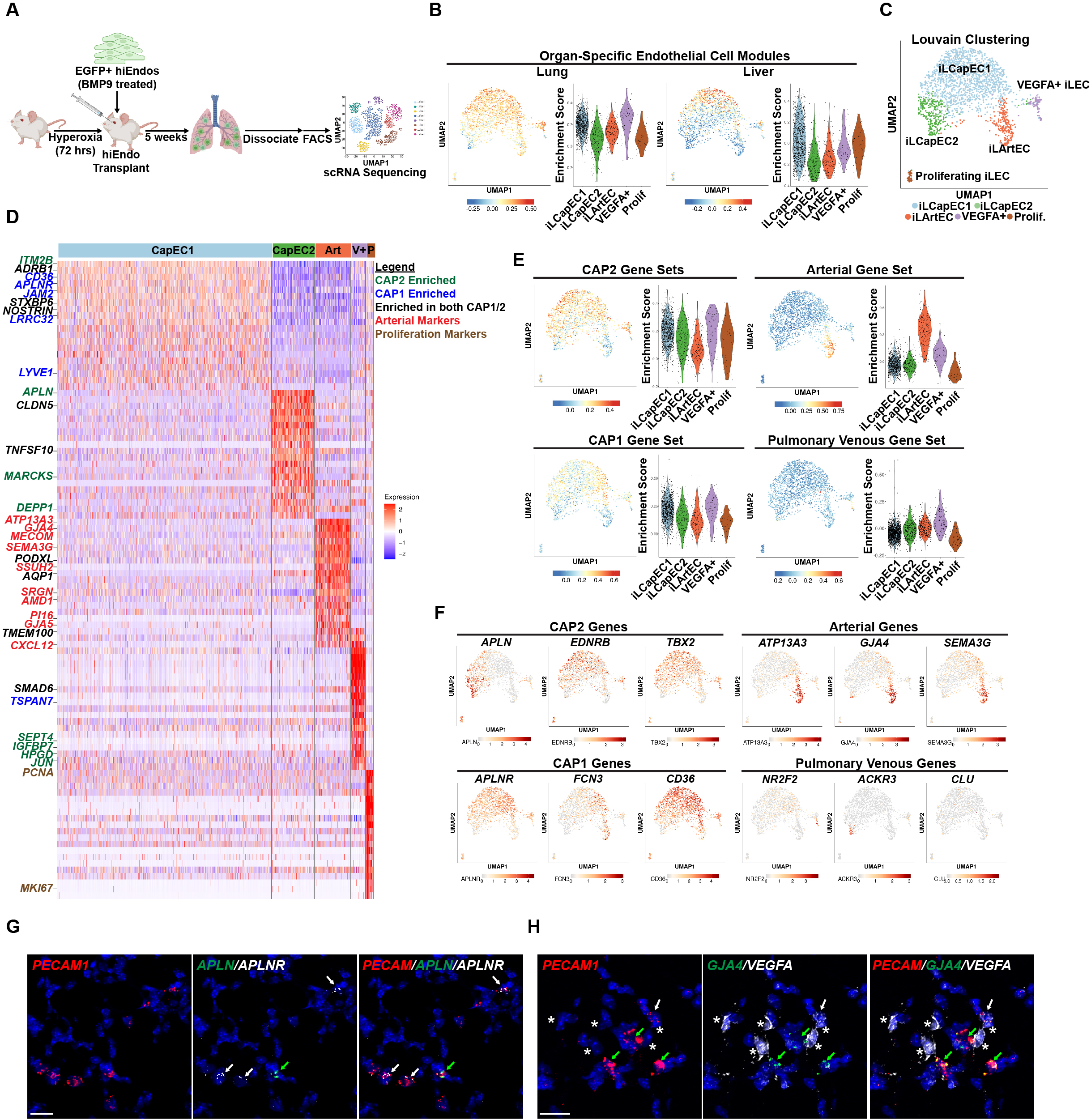
Lung-engrafted hiEndos adopt a lung capillary gene expression profile. (A) Schematic of scRNA-seq profiling of hiEndos 5-weeks post-transplantation into the mouse lung. Data collected from cells pooled from 4 animals. (B) Enrichment scores for the indicated organ-specific endothelial gene sets depicted via UMAP and violin plots. (C) UMAP representation and Louvain clustering (resolution 0.2) of hiEndo transcriptomes 5-weeks post-transplantation. (D) Heatmap representation of top 20 DEGs between post-transplant populations. Genes categories are denoted by color. (E) Enrichment scores for the indicated lung endothelial gene sets^13^ represented by UMAP and violin plots. (F) UMAP plots depicting expression of the indicated gene markers. (G-H) RNA in situ hybridization shows expression of human transcripts including *PECAM1* (red, G-H), *APLN* (green, G), *APLNR* (white, G), *GJA4* (green, H), and *VEGFA* (white, H). N=3 animals per probe set. White arrows indicate *APLNR*+ (G) or *VEGFA*+ (H) hiEndos and green arrows indicate *APLN*+ (G) or *GJA4*+ (H) hiEndos. White asterisks (H) indicate endogenous mouse VEGFA expression. Scale bar (G, H) 20um. Abbreviations: iLCapEC (induced lung capillary endothelial cell), iLArtEC (induced lung arterial endothelial cell), VEGFA+ iLEC (*VEGFA*-expressing induced lung endothelial cell), prolif (proliferating).

### BMPR2 C118W/+ hiEndos exhibit impaired BMP signaling and reduced fitness to engraft in the mouse lung compared to gene-corrected hiEndos

We next leveraged our chimeric human-mouse lung vascular model to explore the mechanism of *BMPR2*-related pulmonary vascular disease, the most common genetic cause of idiopathic and familial PH.^2–5,7^ Towards this end, we generated hiPSCs from two individuals from the landmark 2001 study of a large family kindred with *BMPR2*-related PH.^7^ The hiPSCs from both donors harbored the heterozygous pathogenic *BMPR2* p.(C118W) variant (Figure 5A)^7^, and we used CAS9-mediated gene editing to generate syngeneic *BMPR2* gene-corrected control hiPSCs to be paired with each donor line. All parental and paired corrected lines had normal colony morphology and karyotypes (Figure S5). Both mutant *(BMPR2 C118W/+*) and corrected (*BMPR2 +/+*) hiPSCs were competent for mesodermal and hiEndo differentiation and demonstrated the expected developmental trajectory (Figures S6A and B). We observed a higher efficiency of hiEndo generation on day 5 of differentiation in mutant compared to corrected cells and mutant cells also produced a higher cell yield over serial passages (Figures 5B-D). We assessed the impact of the p.(C118W) variant on BMP signaling and found reduced induction of *ID2* expression in mutant compared to corrected cells on day 2 of directed differentiation after 48 hours of BMP4 exposure (Figures S6A and B). Furthermore, in response to BMP9 treatment, mutant hiEndos showed reduced phospho-SMAD1/5/9 nuclear localization and reduced induction of *ID2* expression (Figures 5E-F). These results demonstrate that *BMPR2 C118W/+* hiEndos exhibit reduced activation of BMP signaling compared to gene-corrected control hiEndos, consistent with prior studies demonstrating a loss-of-function effect of this missense variant.^50,51^

**Figure 5.**
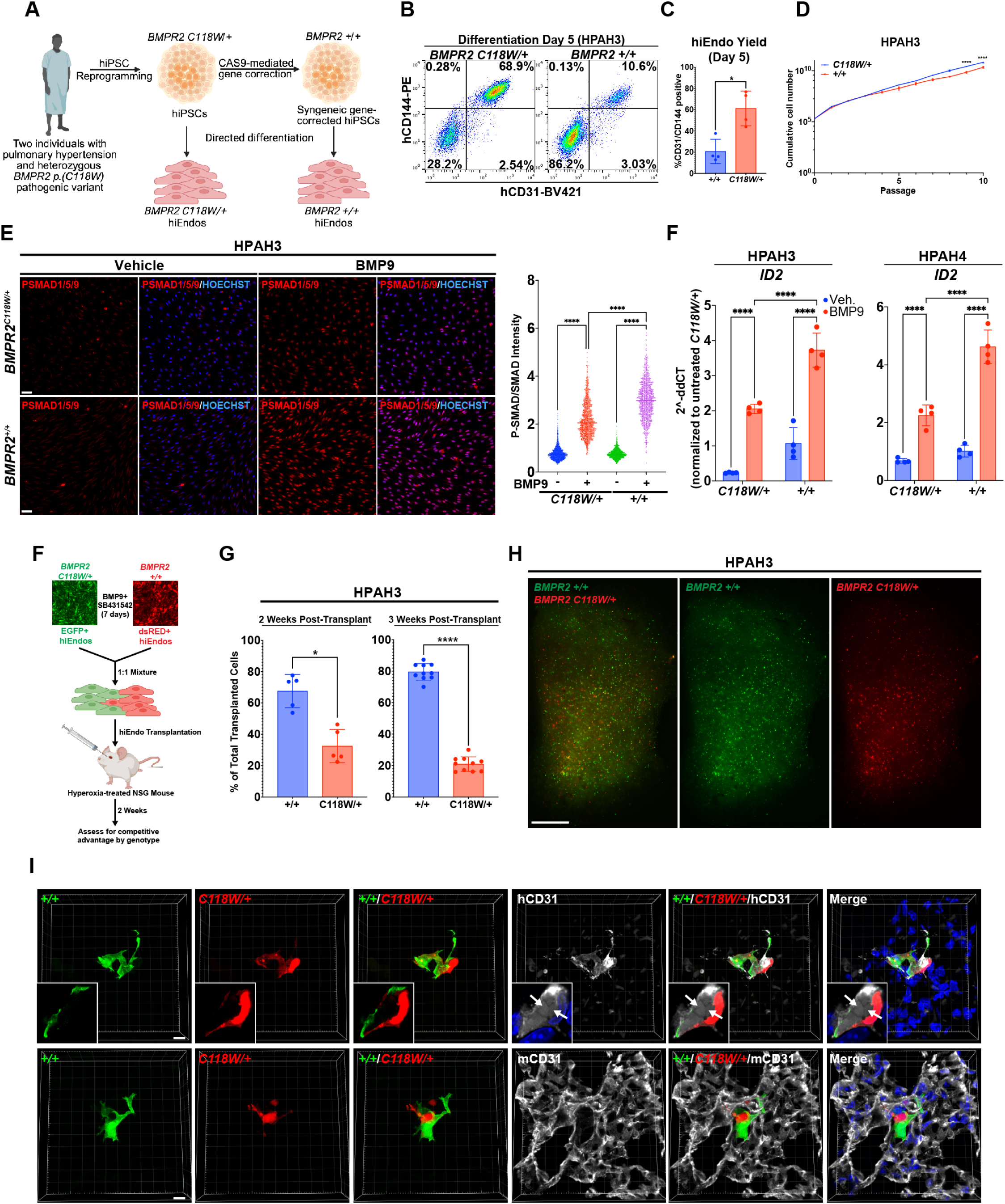
*BMPR2 C118W/+* hiEndos exhibit compromised BMP signaling and reduced engraftment fitness compared to syngeneic gene-corrected cells. (A) Schematic describing the generation of two distinct hiPSC lines from two individuals with pulmonary hypertension and the heterozygous pathogenic *BMPR2 p.(C118W)* variant and syngeneic gene-corrected controls. (B) Flow cytometry for hCD31 and hCD144 on day 5 of differentiation shows increased hiEndo yield in *BMPR2 C118W/+* cells that is quantified in (C; N=4 distinct differentiations per line). (D) Growth assay shows increased cumulative cell yield in *BMPR2 C118W/+* hiEndos compared to gene-corrected controls N=2 experimental replicates of independent wells from the same differentiation, representative data shown from three distinct differentiations. **** p<0.0001 determined by 2-way Anova with Sidak’s multiple comparison test. (E) Immunostaining of hiEndos treated with vehicle or BMP9 for 4 hours and stained with antibodies raised against phospho-SMAD1/5/9 (PSMAD, red). Scale bar 40um. Quantification of nuclear PSMAD fluorescent intensity normalized to total-SMAD1/5/9 intensity is shown to the right. **** p<0.0001 by one-way Anova and Tukey’s multiple comparison test. N=3 experimental replicates from independent wells of the same differentiation, representative data shown from three distinct differentiations. (F) RT-qPCR analysis of *ID2* expression in *BMPR2 C118W/+* and +/+ hiEndos treated with BMP9 for 24 hours. N=4 experimental replicates from independent wells of the same differentiation, representative data shown from three distinct differentiations. **** p<0.0001 by one-way Anova and Tukey’s multiple comparison test. (F) Schematic for competitive transplantation between *BMPR2 C118W/+* and gene-corrected hiEndos. (G) Quantification of the relative contribution of each genotype to the total transplanted hiEndo population 2- and 3-weeks post-transplantation. * p<0.05, **** p<0.0001, Student’s paired two-tailed t-test. Each dot represents an individual animal. (H) Whole-mount fluorescent image of lungs following competitive transplant of *BMPR2 C118W/+* (dsRED) and +/+ (EGFP) hiEndos. Scale bar 2mm. (I) 3D reconstruction of confocal Z stacks of transplanted hiEndos from the competitive lung reconstitution assay showing *BMPR2 C118W/+* (dsRED) and +/+ (EGFP) hiEndos stained with antibodies raised against human CD31 (hCD31, white, top) and mouse CD31 (mCD31, white, bottom). Arrows in insets show red blood cells contained in lumen formed by hiEndos. Experiment replicated with N=3 animals. Scale bar 10um.

We next sought to quantitatively assess the *in vivo* lung endothelial repopulation potential of *C118W/+* compared to +/+ hiEndos using our competitive lung endothelial reconstitution assay. We identified a significant engraftment advantage of +/+ compared to *C118W/+* cells for both HPAH3 and HPAH4 hiEndos, albeit one experiment with HPAH4 hiEndos showed equal engraftment fitness at 2-weeks that diverged at 3-weeks post-transplantation with corrected hiEndos again showing an engraftment advantage (Figures 5F-H, S6C). Engrafted hiEndos from all lines exhibited a similarly complex morphology and maintained human CD31 expression *in vivo*, reproducing results from BU1 hiEndo transplantation studies (Figures 4D and 5I). Red blood cells could be identified within vessel lumens formed from engrafted human endothelia, again confirming the functional connectivity of both mutant and corrected hiEndos to the endogenous microvasculature (Figure 5I insets, arrows).

To determine the impact of the pathogenic *BMPR2* variant on engrafted human lung endothelial cells, we isolated hiEndos from recipient mice for scRNA-seq profiling 3 weeks after competitive lung endothelial reconstitution with *BMPR2 C118W/+* and *+/+* hiEndos (Figure 6A). Moreover, we also included BMP9-treated pre-transplant hiEndos from each hiPSC line to gain a deeper understanding of the transcriptomic changes following hiEndo transplantation (Figure 6A). We first focused on the differences between pre- and post-transplant cells (Figure 6B). hiEndos upregulated their endothelial gene set signatures after transplantation, suggesting further maturation *in vivo* with respect to key endothelial lineage markers (Figure 6C).^13^ We also observed up-regulation of *KLF2*, a marker known to be increased by fluid flow and downregulation of *EDN1*, a marker known to decrease with fluid flow (Figure 6C).^52^ Furthermore, transplantation further augmented expression of a lung-specific endothelial gene set for both normal and mutant iEndos (HPAH3 and HPAH4; Figure 6D). This increased enrichment was not observed for liver or skin gene sets, suggesting lung specificity of *in vivo* hiEndo maturation after orthotopic engraftment in the lung vasculature (Figure 6D). Moreover, hiEndos *in vivo* exhibited higher enrichment of CAP2 and arterial gene sets compared to pre-transplant cells (Figure 6E). Conversely, expression of an ‘early capillary’ gene set, based on published human fetal lung scRNA-seq profiles,^53^ significantly decreased after transplantation (Figure 6E). Taken together, these data indicate endothelial cell maturation and enrichment of a lung endothelial transcriptional program following hiEndo transplantation into the mouse lung.

**Figure 6.**
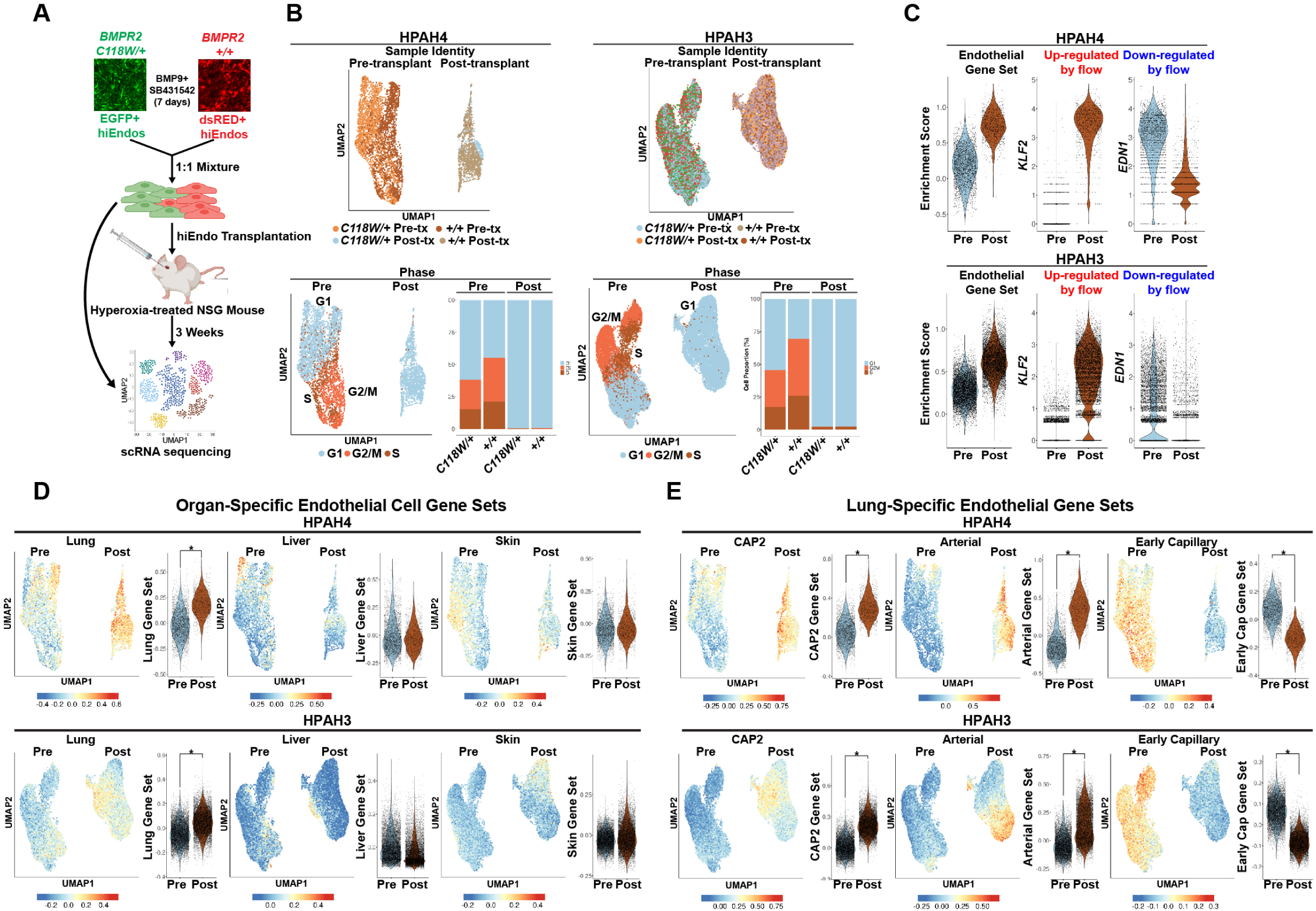
Endothelial maturation and increased lung endothelial gene expression after hiEndo transplantation into the mouse lung. (A) Schematic representation of experiment comparing *BMPR2 C118W/+* and gene-corrected hiEndos both pre- and post-transplant. (B) UMAP representation of scRNA-seq profiles of pre- and post-transplant hiEndos by sample identity (top) and cell cycle phase (bottom). Bar graphs show the relative proportion of cells in each cell cycle phase. (C) Violin plots of pre- and post-transplant hiEndos showing enrichment scores for an endothelial gene set^13^, *KLF2* (up-regulated by flow), and *EDN1* (down-regulated by flow). (D-E) UMAP representation and violin plots showing enrichment scores for organ-specific endothelial gene sets (D)^9^ and lung-specific endothelial gene sets (E)^13^ in pre- and post-transplant cells derived from both HPAP3 and HPAP4 lines. * p<1e-200.

### Novel insights into BMPR2-related PH revealed by patient-derived hiEndos in vitro and in vivo using the chimeric human-mouse lung vascular system

Next, to identify potential mechanisms associated with pathogenesis of *BMPR2*-related PH, we identified differentially expressed transcripts between *BMPR2 C118W/+* and *+/+* hiEndos both before and after transplantation. To determine DEGs in the pre-transplant setting, we generated pre-transplant subplots of the HPAH3/4 datasets (Figure S7A, Tables S7 and S8). We performed gene set enrichment analysis (GSEA) to identify pathways dysregulated in *BMPR2*-mutated cells, and this identified up-regulation of genes involved in the P53 pathway and heme metabolism in the HPAH4 dataset and down regulation of E2F targets, consistent with decreased proliferation, in both HPAH3 and HPAH4 pre-transplant datasets (less G2/M and S phase; Figure 6B; GSEA Figure S7B, Tables S9 and S10). Although, there were no statistically significantly up-regulated Hallmark pathways in the HPAH3 pre-transplant dataset (Table S10), we sought to determine significantly up- and down-regulated individual genes in *C118W/+* vs +/+ hiEndos that are common to both pre-transplant datasets. We identified 127 shared up-regulated and 307 shared down-regulated genes (Figure S7C, Tables S7 and S8), with the long non-coding RNA *H19* as one of the top up-regulated genes in mutated pre-transplant hiEndos in both data sets (Figure S7D). Genes involved in the oxidative stress response were also commonly up-regulated in *BMPR2*-mutated cells in the pre-transplant setting (Figure S7E). With respect to down-regulated genes, we identified reduced *BMPR2* expression in *C118W/+* compared to +/+ cells, similar to prior studies of *BMPR2*-mutated hiEndos *in vitro* (Figure S7F).^23^ We also identified down-regulation of the negative feedback inhibitor and direct target of the BMP pathway, *SMAD6* (Figure S7F).^54,55^

Finally, to define the disease signature of engrafted cells, we subclustered just the post-transplant hiEndos (Figure 7A, Table S11 and S12) and assigned the same annotations revealed by our BU1 post-transplant dataset (Figure 4), demonstrating the reproducibility of this system across hiPSC lines and across experiments. To identify the transcriptional consequences of the p.(C118W) variant, we performed differential gene expression analysis in post-transplant cells by genotype in both datasets and identified 31 commonly up-regulated genes and 36 commonly down-regulated genes in *BMPR2 C118W/+* compared to +/+ hiEndos (Figure 7C, Table S13 and S14). This analysis again identified the lncRNA *H19* as the top DEG upregulated in variant cells in both datasets, but with increased expression and greater difference between variant and corrected cells following lung engraftment (Figure 7D and E). We validated this result by RNA *in situ* hybridization showing *H19* expression exclusively in *C118W/+* cells in vivo (Figure S7G). We also identified up-regulation of the *AHR* receptor and its downstream target *CYP1B1* in *C118W/+* cells in both HPAH3 and HPAH4 datasets (Figure 7D and F), and upregulation of an additional AHR downstream target, *TIPARP*, in the HPAH3 dataset (Figure 7F). This analysis also identified up-regulation of genes in *C118W/+* cells related to TGF-beta signaling (*TGFBR2* and *SKIL*); matricellular components (*FN1*, *SPARC*); signaling ligands (*JAG1*, *NRG3*); transcription factors (*CHCHD2*, *CTNNB1*, *TBX3*, etc.); and others (Figure 7F).

**Figure 7.**
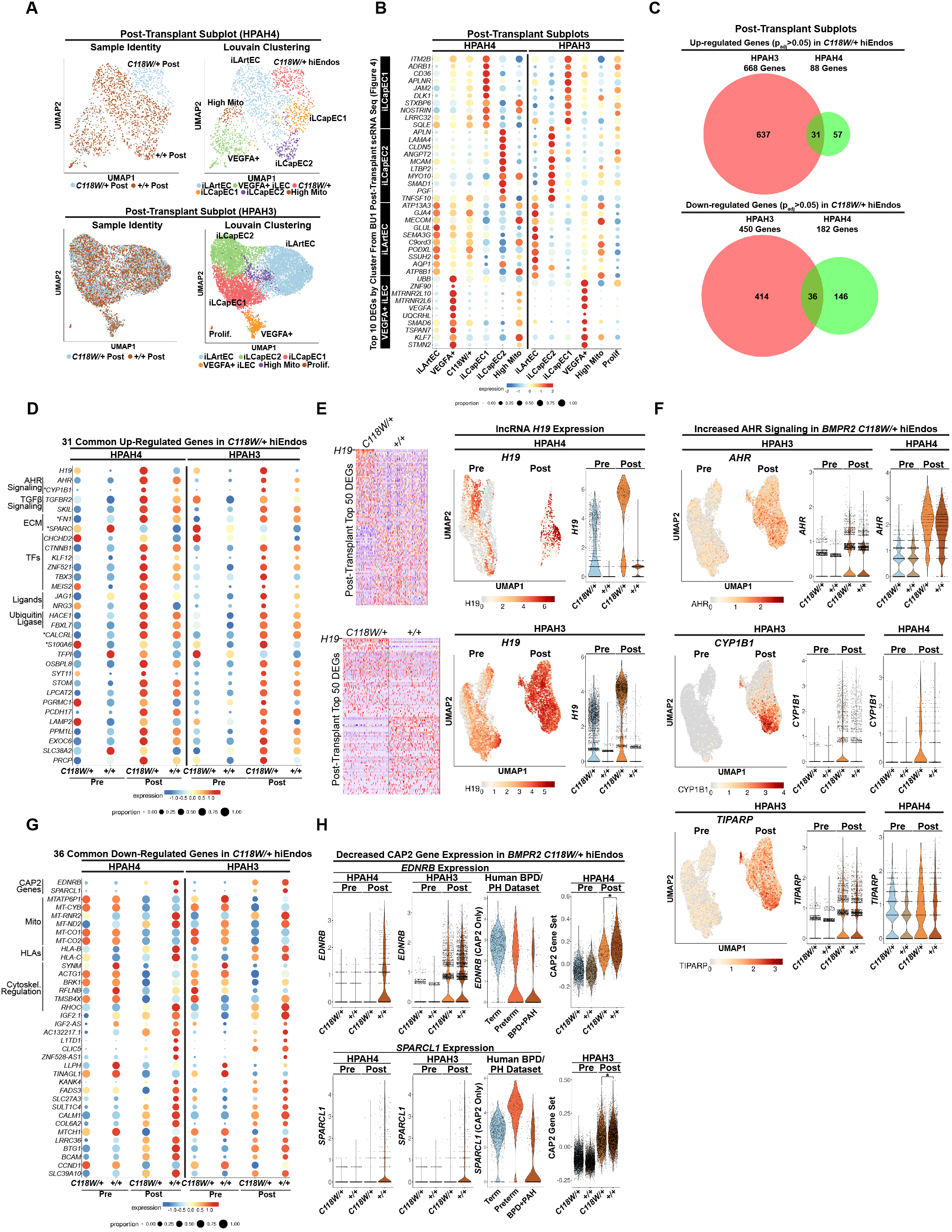
scRNA-seq reveals defects in capillary gene expression, increased lncRNA H19 expression, and elevated AHR signaling in post-transplant *BMPR2 C118W/+* hiEndos. (A) UMAP representation of post-transplant subplots of HPAH3 and HPAH4 hiEndos from Figure 6 showing sample identity and Louvain clustering. (B) Dot plots from the HPAH3/4 post-transplant data sets demonstrates expression of the top 10 DEGs between cell clusters defined from scRNA-seq from Figure 4. (C) Venn diagram showing overlap of genes that are significantly up- or down-regulated (padj < 0.05) in *BMPR2 C118W/+* post-transplant hiEndos compared to gene-corrected controls. (D) Dot plot reveals expression of the 31 common up-regulated genes in pre- and post-transplant hiEndos by genotype. *Denotes genes identified in other pulmonary hypertension models and human patient samples via scRNA-seq. (E) Heat maps showing that *H19* is the top up-regulated DEG in both post-transplant datasets. *H19* expression in pre- and post-transplant cells is revealed via UMAP representation and violin plots of pre- and post-transplant cells by genotype. (F) Expression of AHR, CYP1B1, and TIPARP depicted by UMAP representation and violin plots of pre- and post-transplant hiEndos by genotype. (G) Dot plot reveals expression of the 36 common down-regulated genes in pre- and post-transplant hiEndos by genotype. (H) Violin plots show expression of *EDNRB* and *SPARCL1* in HPAH3/4 pre- and post-transplant hiEndos and in CAP2 cells from a human BPD/PH scRNA-seq dataset^56^ in addition to enrichment scores for a CAP2 gene set.^13^ * p<1e-10. Abbreviations: mito (mitochondrial), BPD (bronchopulmonary dysplasia), PH (pulmonary hypertension), TFs (transcription factors).

With respect to down-regulated genes, *BMPR2* mutated hiEndos exhibited decreased expression of the CAP2 markers *EDNRB* and *SPARCL1* that is only appreciated in the post-transplant setting (Figure 7G and H). To investigate further a capillary gene expression defect in *BMPR2 C118W/+* cells, we found decreased enrichment of a CAP2 gene set in *C118W/+* post-transplant hiEndos, implicating a defect in lung-specific capillary gene expression resulting from the pathogenic *BMPR2* variant (Figure 7E). This phenomenon was also observed in a previously published dataset with decreased expression of *EDNRB* and *SPARCL1* in CAP2 cells in infants with bronchopulmonary dysplasia and PH compared to term and preterm infants (Figure 7H).^56^

To determine the context-dependent nature of our transcriptomic analyses, we analyzed the overlap between the shared up- and down-regulated genes between the pre- and post-transplant datasets. This identified only 4 common up-regulated (*H19, CHCHD2, TGFBR2, S100A6*) and 2 common down-regulated genes (*LLPH, SYNM*) that were shared in both pre- and post-transplant datasets (Figure S7H). Overall, these data reveal cellular derangements resulting from the *BMPR2* p.(C118W) missense variant *in vivo* including diminished functional lung endothelial reconstitution capacity, increased lncRNA *H19* expression, increased AHR signaling, and impaired lung capillary gene expression. Moreover, these changes are largely context-dependent with most only revealed in the *in vivo* lung-engrafted context.

## Discussion

Our results indicate that disease-specific hiPSCs can be harnessed for *in vitro* lung endothelial pre-patterning into lung-specific endothelia that facilitates *in vivo* functional engraftment of the resulting human endothelial cells, generating an *in vivo* chimeric human-mouse lung vascular model with augmented lung-specific transcriptomic expression and amplified disease-modeling potential. To develop this approach, we defined a novel directed differentiation and expansion protocol to generate billions of hiEndos *in vitro*. One key challenge with respect to cultured endothelial cells is the propensity of cells to transition towards a fibroblast-like state as well as losing their proliferative potential over serial passages.^57^ Our results show that prolonged maintenance of cell fate and proliferative potential through inhibition of both TGF-beta and Notch signaling can generate billions of cells in several weeks from a single differentiation. This strategy overcomes a key hurdle in generating sufficient cell quantity for transplantation studies that may be applied to the development of cell-based therapies for pulmonary vascular diseases.

Developing a competitive lung endothelial reconstitution assay provided a key quantitative functional readout for testing whether disease-associated mutations, such as in *BMPR2*, or pre-treatment of human cells with BMP signaling stimulation impacts either their endothelial reconstitution potential or their molecular phenotypes when studied *in vivo* in the environment of the same lung. We found that BMP9 treatment of hiEndos prior to transplantation promotes lung engraftment. A similar strategy also encouraged transplantation of mouse iPSC-derived endothelial cells in neonatal mice.^58^ BMP9 is a key signaling molecule for lung vascular development and pathogenic variants in several components of this signaling pathway including the ligand (*GDF2*), receptors (*ACVRL1*, *BMPR2*), and other components (*ENG*) are associated with pulmonary hypertension in humans.^2,5,7,59–66^ Our data demonstrate that hiEndos are competent to respond to BMP9 and up-regulate the lung-specific endothelial marker *TMEM100* and other lung capillary programs *in vitro*. We reasoned a ‘like-into-like’ model wherein patterning of hiEndos towards a lung-specific molecular profile promotes their engraftment into the lung microvasculature and similar organotypic patterning strategies may be needed for endothelial reconstitution in other organ systems.

We found our novel chimeric human-mouse lung vascular model significantly extends the already powerful disease modeling potential of hiPSC-derived endothelial lineages. When partnered with *in vitro* disease modeling, hiEndos after *in vivo* engraftment may reveal disease-associated programs that emerge or augment following orthotopic transplantation into a context that more fully includes the multitude of cell types and environmental variables present in an air-breathing, vascularized organ *in vivo*. We note for example, that transplanted hiEndos are localized to their native tissue context and experience the endogenous niche cues and inter-organ crosstalk required for lung endothelial identity. Lung-engrafted hiEndos also experience tissue-specific biophysical forces such as fluid flow and stretch from breathing that are essential to the development of lung cell fates.^67^ Our transplantation model generates expanded cellular diversity with both capillary- and arterial-like populations enabling studies of the impact of pathogenic gene variation on multiple cell types in a single experiment. This *in vivo* approach along with emerging *in vitro* vascularized lung organoid models reflects significant recent advances in modeling human lung vascular development and disease.^68^

We applied our chimeric human-mouse lung vascular system to model genetic pulmonary vascular disease by co-transplantation of hiEndos derived from patients with *BMPR2*-related PH along with syngeneic gene-corrected control cells. Using the quantitative competitive lung reconstitution assay, we observed an engraftment advantage of gene corrected compared to *BMPR2* mutated hiEndos. Critical to the success of cellular replacement therapies is the ability of gene-corrected cells to expand and out-compete their diseased counterparts. Our data suggests that gene-corrected hiEndos may be poised to expand following transplantation into a diseased context to correct vasculopathy. Through scRNA-seq, we also show several derangements downstream of the *BMPR2* mutation that have not been previously implicated in disease pathogenesis. Of note, many of these derangements are only observed following hiEndo lung engraftment, demonstrating the utility of this model to gain insights into PH pathogenesis. Importantly, we have narrowed down potential candidate pathways through analysis of cells generated from two distinct individuals with *BMPR2*-related PH with the same heterozygous p.(C118W) variant. This includes up-regulation of the lncRNA *H19*, which has recently been shown to be up-regulated in human idiopathic PH lungs, and *H19* knockout prevented pulmonary hypertension in a rat hypoxia model implicating a key role in disease pathogenesis^69^. Up-regulation of *H19* has been shown to be critical in the pathogenesis of heart failure secondary to pulmonary hypertension and *H19* suppression rescued right ventricular failure in a monocrotaline-induced PH rat model.^70^ Future studies are now needed to address the mechanism and consequences of *H19* up-regulation in a diseased context.

We also found evidence of increased AHR signaling in engrafted BMPR2 variant hiEndos including up-regulation of the *AHR* receptor as well as its downstream target *CYP1B1*. Interestingly, a recent study showed that deficiency of AHR in rats was protective against PH development in the SU5416/hypoxia model.^71^ In fact, SU5416 activation of AHR signaling rather than VEGF inhibition was determined to be causative for PH in this model system, demonstrating a key role for AHR signaling as a driver of disease pathogenesis.^71^ Prior studies identified a correlation between reduced *CYP1B1* expression and PH development in individuals with *BMPR2*-related PH. Our study and several other scRNA-seq studies of human and rat PH show increased *CYP1B1* expression in diseased endothelial cells, implicating a more complex role for *CYP1B1* in disease pathogenesis.^72–76^ Additional genes of interest identified as upregulated in our study include *FN1*, *SPARC*, *CALCRL*, and *S100A6* were also increased in scRNA-seq studies of human and murine PH models, although the mechanistic contribution to disease pathogenesis is unknown.^56,72–77^

Our transplantation model also shows a deficiency in CAP2 gene expression resulting from the pathogenic *BMPR2* variant. This includes down-regulation of the CAP2 markers *EDNRB* and *SPARCL1* and reduced enrichment scores of a CAP2 gene set. This phenomenon has also been observed in a mouse model of *Bmpr2*-related PH and several human PH scRNA-seq datasets show reduced CAP2 cells with abnormal gene expression in diseased lungs.^56,74,78^ CAP2 cells generate the majority of the surface area of the lung capillary bed and rarefaction or dysfunction of this population may lead to elevated pulmonary pressures over time.^12,79,80^ This reduction in CAP2 gene expression is also seen in scRNA-seq studies of human PH samples. Future studies possibly utilizing this novel system can now focus on identifying convergent and distinct mechanisms of disease in other forms of genetic PH.

### Limitations

One limitation of our system is the mixed CAP1 and CAP2 gene expression profiles expressed in many of the engrafted hiEndos.^68^ This may reflect the intermediate capillary cell state recently noted during development and in response to lung injury.^81^ Of note, this does not reflect the recently defined pathologic transitional state given the lack of *NTRK2* expression.^82^ While the identified iLArtEC populations do not reside in large vessels, this may reflect capillary endothelium that reside on the arterial side of the vascular bed as observed in the lung and other organ systems.^83–85^ Our transplantation model also generates hiEndos with high *VEGFA* expression reminiscent of autocrine signaling observed in endothelial cells during injury repair following influenza infection.^86^ These hiEndos may be responding to regions of ongoing repair following hyperoxia injury. Studying how each particular hiEndo-derived endothelial subtype functions or further matures with extended time *in vivo* or after various mouse lung injury exposures should help with interpretation of the various molecular phenotypes present in hiEndos after engraftment. We also note a limited reconstitution of the lung endothelium with engrafted human cells (∼1 to 2% at 2-weeks). Additional strategies to enhance cellular engraftment are needed to alter vascular physiology or rescue disease phenotype to test the considerable therapeutic potential of hiEndos to reconstitute the pulmonary vascular endothelium as a cell-based therapy in settings where dysfunctional or damaged pulmonary endothelium drives disease.

---

**Figure S1.**
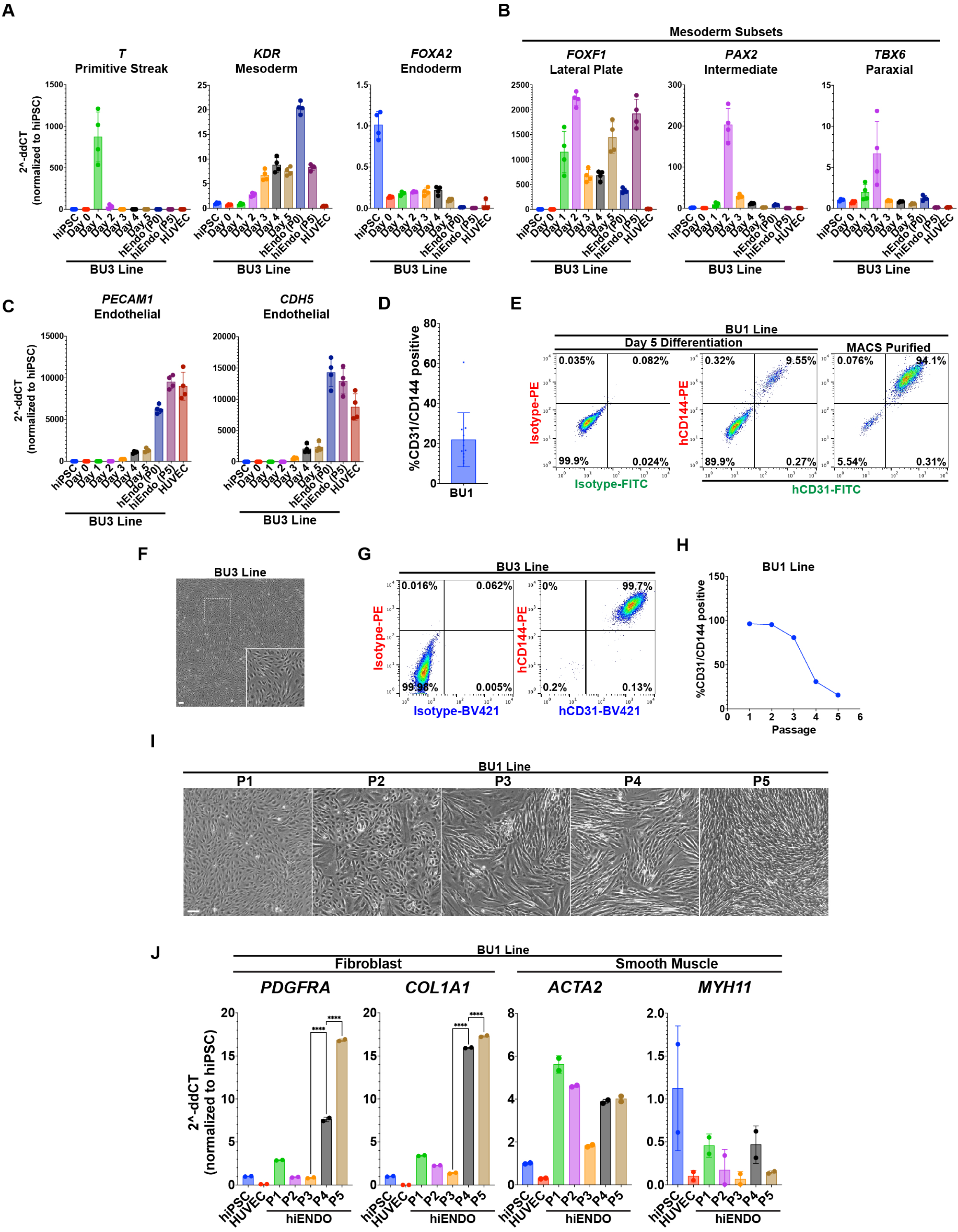
Additional characterization of hiEndo differentiation and maintenance. (A-C) Fold change in the expression of indicated transcripts in hiPSC-derived cells (BU3 clone) on each day of hiEndo differentiation, in purified hiEndos at P0 and P5, and in HUVECs (RT-qPCR). N=4 experimental replicates from independent wells of the same differentiation. (D) Quantitation of hiEndo differentiation efficiency on day 5 of differentiation across 12 distinct differentiations. (E) Flow cytometry of human CD31, human CD144, and isotype controls pre- and post-MACs purification of hiEndos on day 5 of the differentiation. N=1. (F) Phase contrast microscopy of BU3 hiEndos showing cobblestone morphology. Scale bar 50um. (G) Flow cytometry of human CD31, human CD44, and isotype controls in BU3 hiEndos, the plot is representative of 4 experimental replicates of independent wells from the same differentiation. (H) Quantification of the percent of CD31/CD144 double positive hiEndos by flow cytometry at each passage. N=1 differentiation (BU1). (I) Phase contrast microscopy of BU1 hiEndos at sequential passages. Scale bar 50um. Results are representative of 2 experimental replicates of independent wells from the same differentiation. (J) RT-qPCR analysis of fold change in transcript expression of markers of fibroblast and smooth muscle identity in hiEndos at sequential passages. N=2 experimental replicates of independent wells from the same differentiation. **** p<0.0001 by one-way Anova with Tukey’s multiple comparisons.

**Figure S2.**
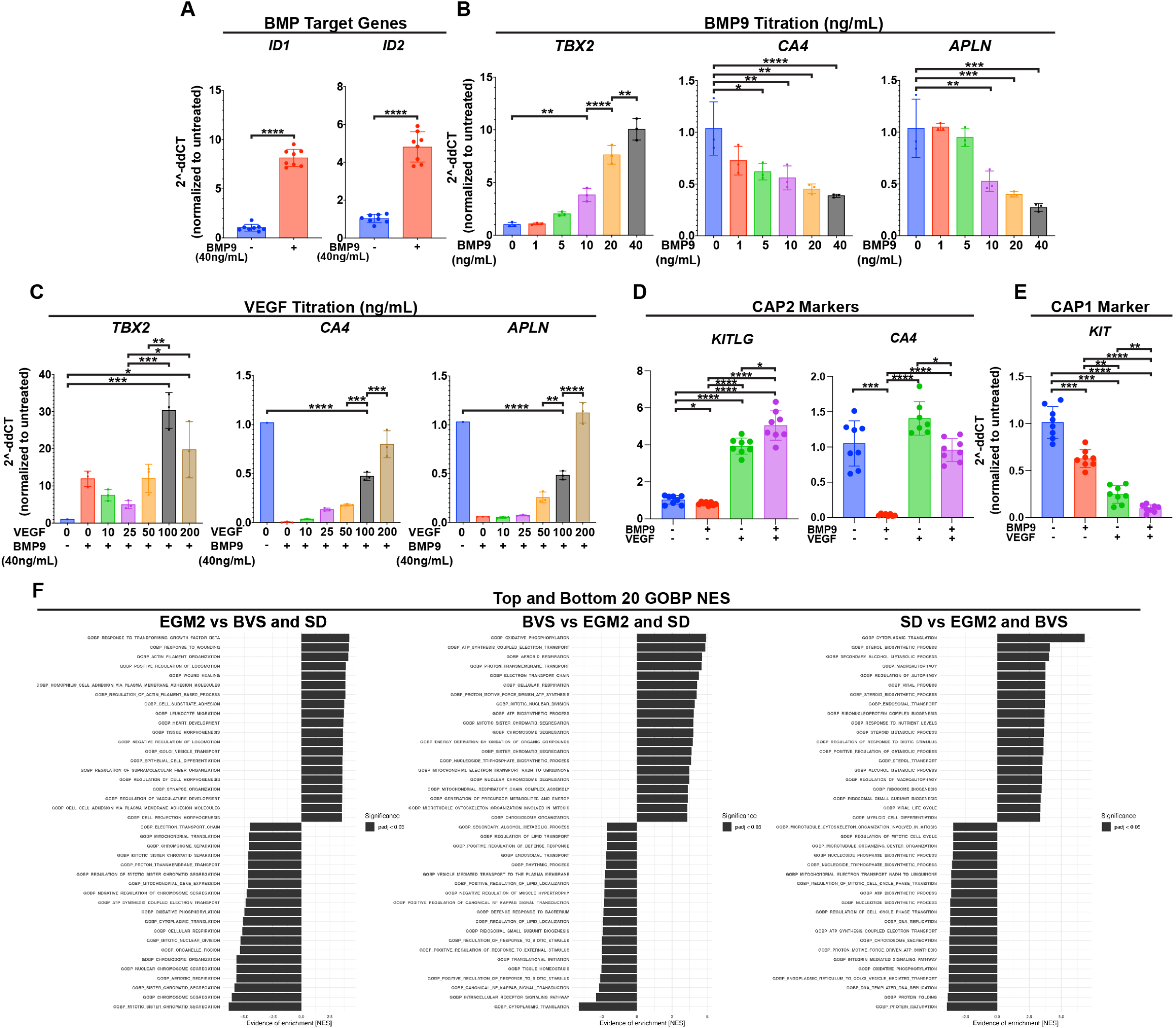
Expression of CAP2 and CAP1 genes in hiEndos treated with BMP9 and VEGFA. (A) Fold change in expression (RT-qPCR) of the indicated transcripts in BU1 hiEndos treated with vehicle or BMP9 for 7 days. **** p<0.0001 by two-tailed Student’s paired t-test. N=8 biological replicates consisting of 8 distinct differentiations. (B) As in (A) except using varying concentrations of BMP9 for 7 days. N=3 experimental replicates from independent wells of the same differentiation, representative data is shown from three distinct differentiations. * p<0.05, ** p<0.01, *** p<0.001, **** p<0.0001 by one-way Anova and Tukey’s multiple comparisons. (C) as in (B) except using varying concentrations of VEGFA in the presence of BMP9. N=3 experimental replicates from independent wells of the same differentiation, representative data is shown from three distinct differentiations. * p<0.05, ** p<0.01, *** p<0.001, **** p<0.0001 by one-way Anova and Tukey’s multiple comparisons. (D and E) As in (A) comparing hiEndos treated alone or in combination with BMP9 (40ng/mL) and VEGF (200ng/mL). N=8 biological replicates. * p<0.05, ** p<0.01, *** p<0.001, **** p<0.0001 by repeat measurement one-way Anova with Tukey’s multiple comparisons. GSEA analysis (GOBP) from the scRNA-seq dataset (Figure 2) comparing each media condition against the other 2 conditions. Black bars indicate significant differences p_adj_<0.05; data are further detailed in supplemental tables S3-5.

**Figure S3.**
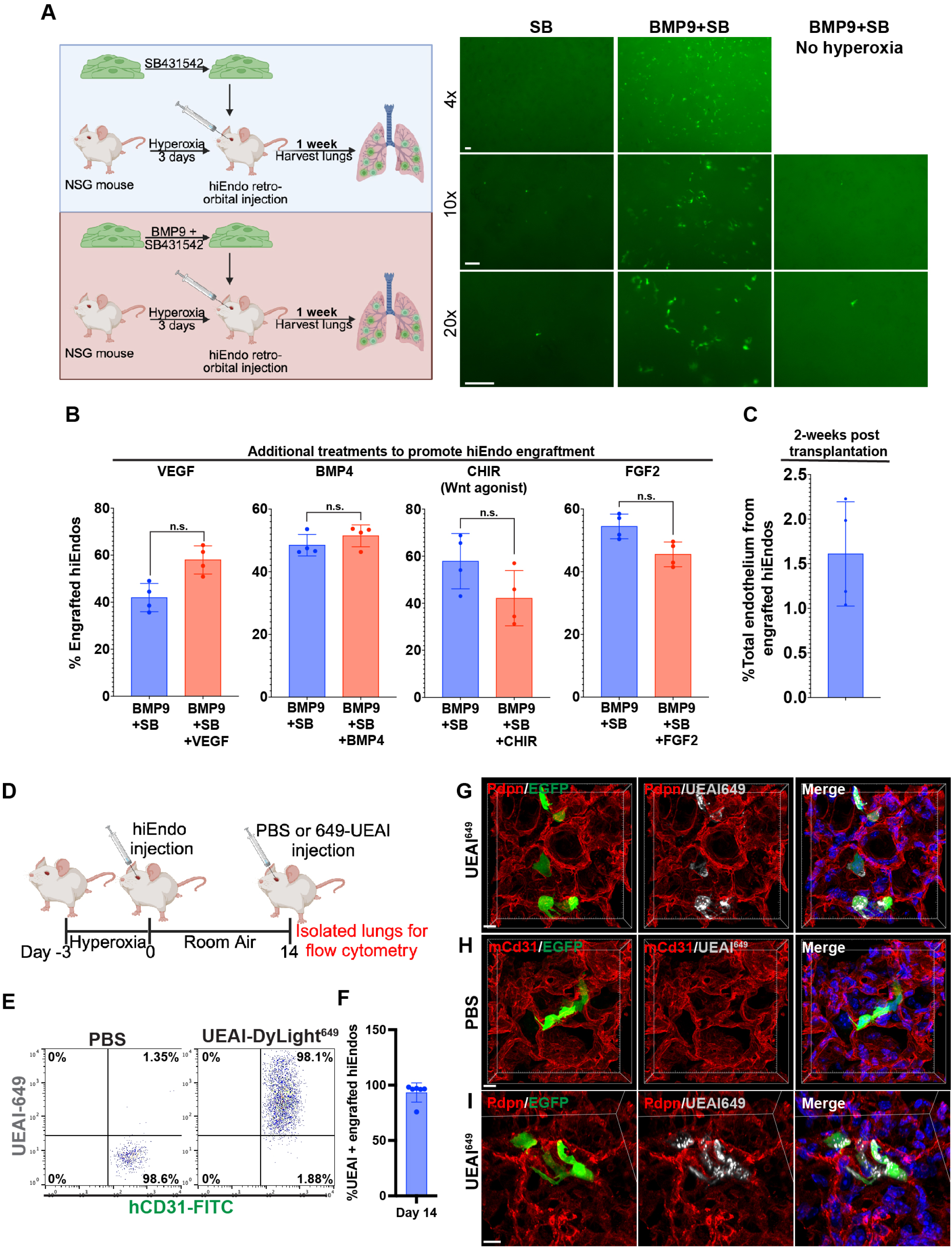
Additional analysis of hiEndo transplantation. (A) Schematic of transplantation studies where SB431542-treated and BMP9+SB431542-treated hiEndos are injected into different mice and EGFP+ signal is used to determine transplantation efficiency (left). Whole mount fluorescent images of freshly dissected lungs that were transplanted with hiEndos treated with SB41542 (SB) or SB+BMP9 for 7-days prior to injection and analyzed 1 week following transplantation (right). N=2 per condition. The far right panel reflects animals that did not undergo hyperoxia and were then transplanted with SB+BMP9-treated hiEndos. N=6 animals. Scale bar 50um. (B) Quantitation of competitive transplantation assay of hiEndos treated with the indicated growth factors and the relative proportion of each cell population quantified by flow cytometry for EGFP+ and dsRED+ hiEndos. N=4 animals per condition. N.s.= not significant by Student’s paired t-test. (C) Quantitation of the percent reconstitution of the mouse lung endothelium with hiEndos 2-weeks post-transplantation measured by flow cytometry. N=4 animals. (D) Schematic of intravascular labeling experiment to assess for connectivity of hiEndos to the endogenous mouse vasculature. (E) Flow cytometric analysis of transplanted hiEndos following intravascular injection of PBS or UEAI-DyLight-649 and immunostaining for hCD31. (F) Quantification of intravascularly labeled transplanted hiEndos by flow cytometry across 6 animals. (G-I) 3D reconstruction of confocal Z-stacks of EGFP+ hiEndos 2-weeks following transplantation. Tissue sections are immunostained for Pdpn (G and I, red) and mCd31 (H, red). Animals were injected with either PBS or UEAI-DyLight-649 (white) prior to dissection. Hoechst labels nuclei (blue). Scale bar (G-I) 10um. Abbreviations: CHIR99021 (CHIR), Ulex europaeus agglutinin I (UEAI).

**Figure S4.**
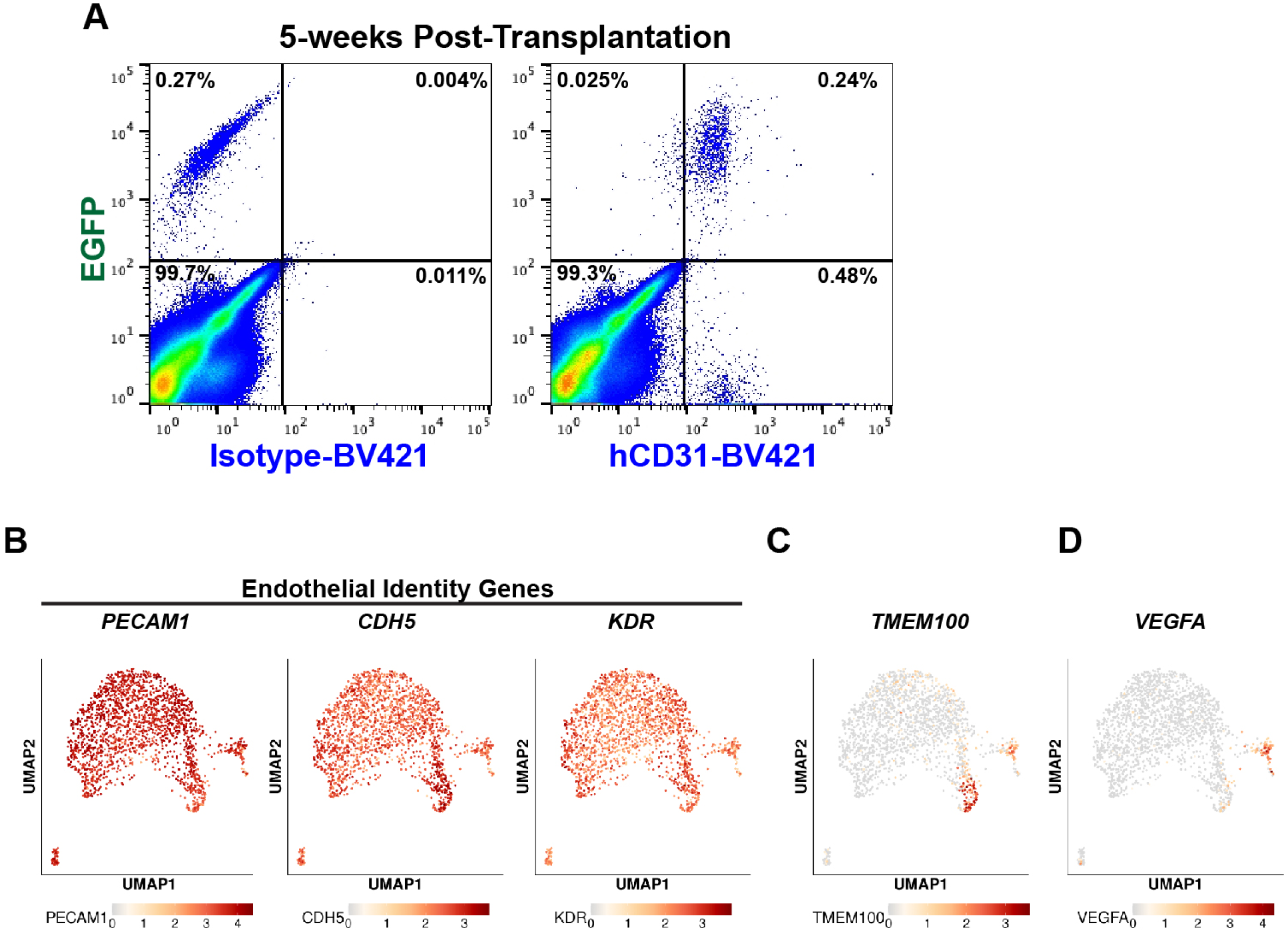
Gene expression analysis of hiEndos 5-weeks post-transplantation in the mouse lung by scRNA-seq. (A) Flow cytometric analysis of mouse lung single cell digests from animals 5-weeks post-transplantation with EGFP-labeled, BMP9+SB-treated hiEndos and stained with either an isotype or human CD31 BV421-conjugated antibodies. (B) UMAP representation of post-transplant BU1 hiEndos showing expression of key endothelial lineage markers. (C and D) As in (B) except showing expression of *TMEM100* (C) or *VEGFA* (D) expression in post-transplant hiEndos.

**Figure S5.**
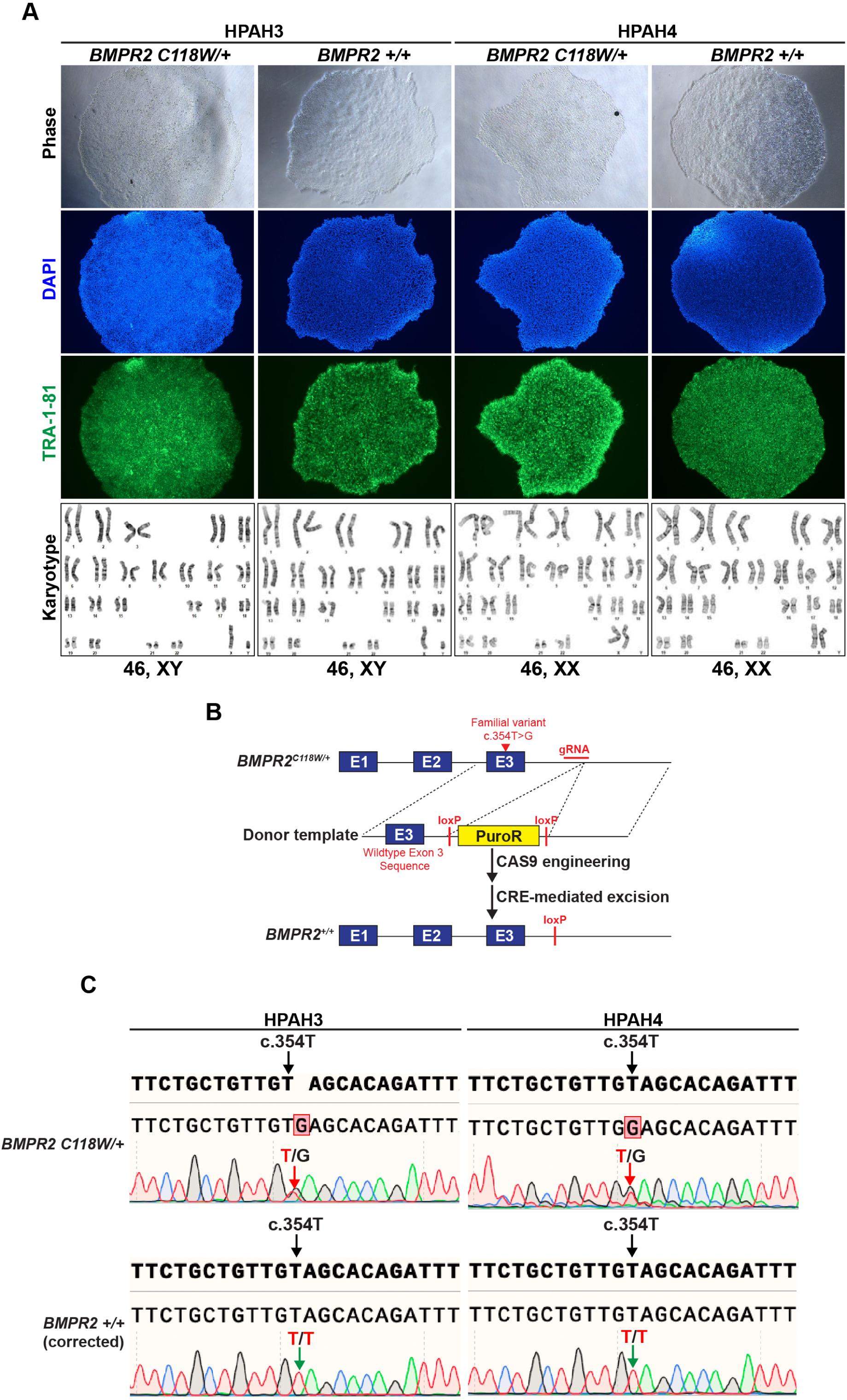
Characterization of iPSCs and CAS9-mediated gene editing to correct the *BMPR2* p.(C118W) variant. (A) Phase contrast (top), DAPI staining (2^nd^ row), TRA-1-81 immunofluoresence (3^rd^ row), and karyotypes of the HPAH3 and HPAH4 hiPSC lines as well as the syngeneic CAS9-corrected lines. (B) Schematic of strategy to correct the *BMPR2* p.(C118W) variant via CAS9-mediated gene editing. Note the presence of a loxP scar after cre-mediated recombination to excise the floxed puromycin resistance cassette (PuroR). (C) Chromatograms from Sanger sequencing showing the heterozygous c.354T>G variant in HPAH3 and HPAH4 lines in addition to the successful gene correction in the edited lines.

**Figure S6.**
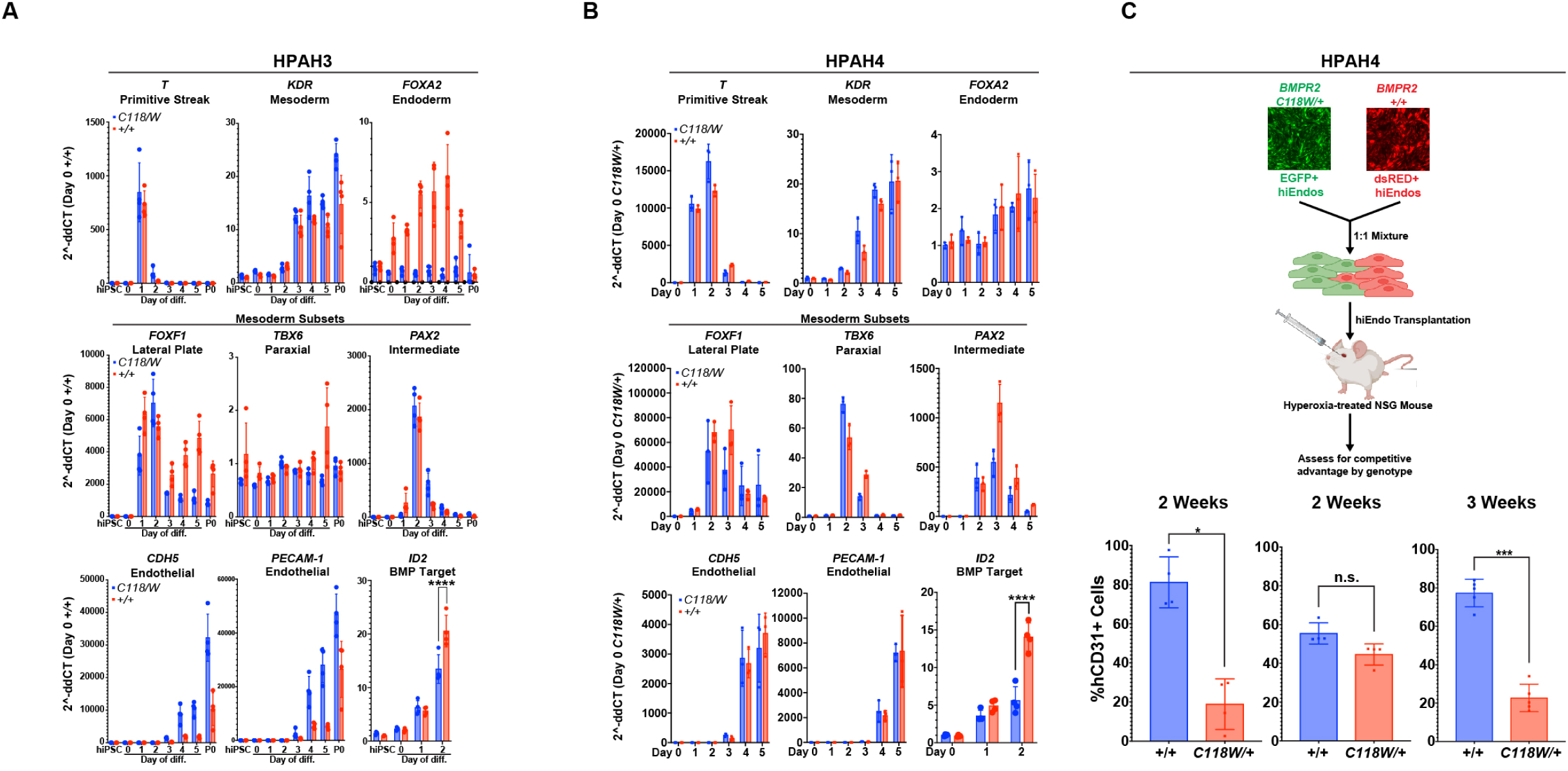
Additional analysis of HPAH3 and HPAH4 hiEndos. (A) RT-qPCR analysis of fold change in expression of the indicated genes in cells from each day of the hiEndo directed differentiation protocol in addition to purified P0 hiEndos. hiPSCs are included as a control. N=4 experimental replicates. **** p<0.0001 by two-way Anova with Tukey’s multiple comparisons. (B) As in (A) except using the HPAH4 line. (C) (top) Schematic of competitive transplantation assay using HPAH4 *BMPR2 C118W/+* vs syngeneic gene-corrected hiEndos. Cells were analyzed at the indicated time points. N=4 animals for the 2-week studies and N=5 for the 3-week time point. * p<0.05, *** p<0.001, n.s. not significant by Student’s two-tailed paired t-test.

**Figure S7.**
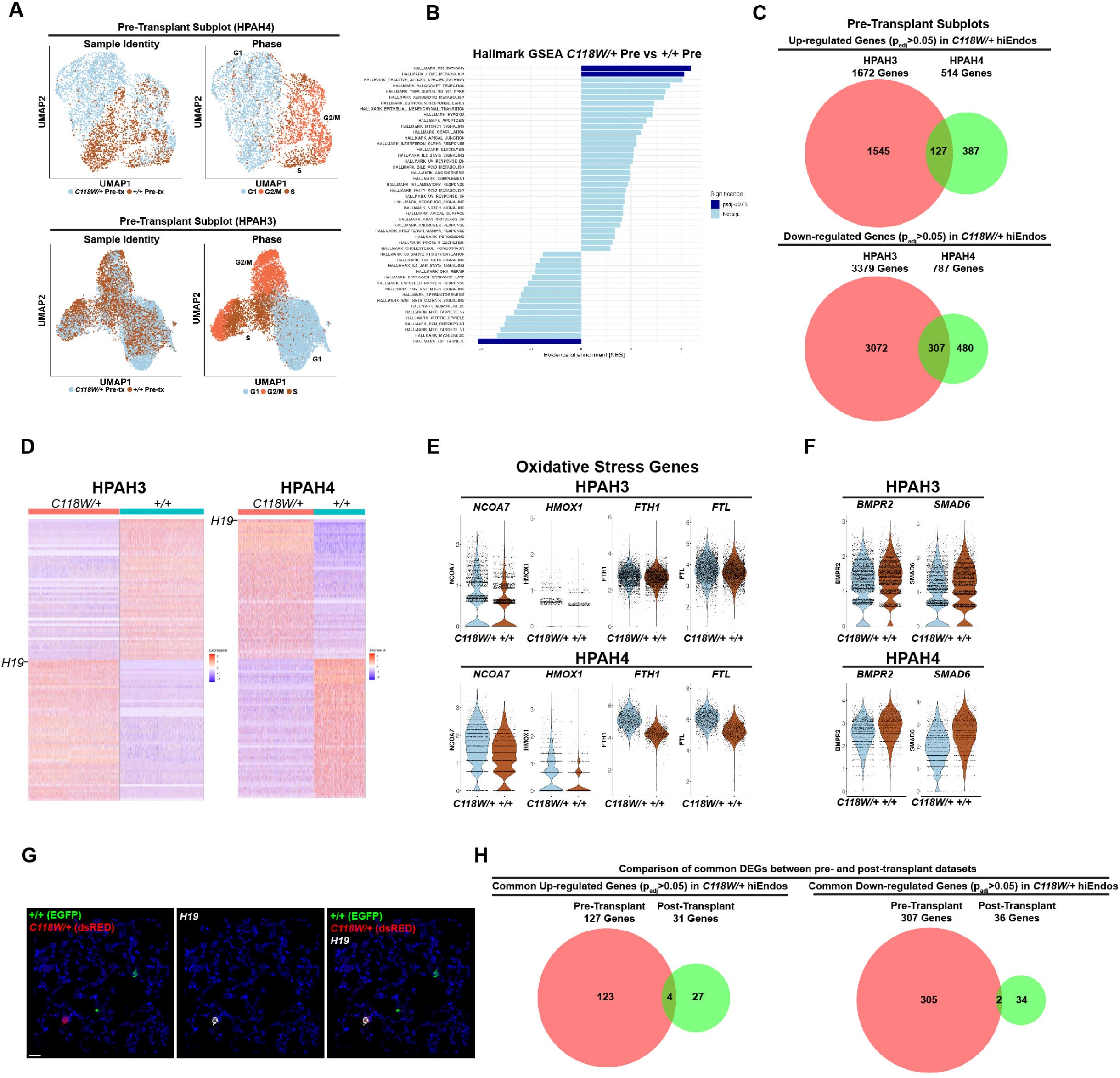
Analysis of HPAH3 and HPAH4 pre-transplant hiEndos by scRNA-seq. (A) UMAP representation of HPAH3 and HPAH4 pre-transplant subplots showing Sample Identity and cell cycle phase. (B) Hallmark GSEA analysis of *BMPR2 C118W/+* vs *+/+* pre-transplant hiEndos. Dark blue lines reflect padj<0.05. (C) Venn diagram showing overlap of genes that are significantly up- or down-regulated (padj < 0.05) in *BMPR2 C118W/+* pre-transplant hiEndos compared to gene-corrected controls. (D) Heatmaps with row-normalized z scores for the top 50 DEGs by genotype in pre-transplant HPAH3 and HPAH4 hiEndos, showing *H19* as the top up-regulated gene in both datasets. (E) Violin plots demonstrating key up-regulated genes in C118W/+ pre-transplant hiEndos related to oxidative stress. (F) As in (E) showing key down-regulated genes in C118W/+ pre-transplant hiEndos including the *BMPR2* receptor and *SMAD6*. (G) RNA in situ hybridization using RNAscope to detect expression of *EGFP*, *dsRED*, and *H19* transcripts in animals undergoing competitive transplant with the HPAH3 line 3-weeks post-transplantation. Note that *H19* transcripts are exclusively identified in mutant cells as determined by fluorophore detection. Microscopy results are representative of N=3 animals. (H) Venn diagrams showing the overlap of significantly up- or down-regulated genes (padj < 0.05) between pre- and post-transplant datasets by genotype.

## Materials and Methods

### Contact for Reagents and Resource Sharing

All unique/stable reagents generated in this study are available from the corresponding author with a completed Materials Transfer Agreement. Pluripotent stem cell lines generated in this study are available from the CReM Biobank at Boston University and Boston Medical Center and can be found at crem.bu.edu. Further information and requests for reagents may be directed to, and will be fulfilled by, the corresponding author Darrell Kotton.

### Mice

NSG mice (NOD.Cg-Prkdc^scid^ Il2rg^tm1Wjl^/zJ) were obtained from The Jackson Laboratory (strain # 005557). Animal methods were approved by the Institutional Animal Care and Use Committee of Boston University School of Medicine and husbandry was performed in facilities overseen by the Animal Science Center at Boston University. The transplantation experiments were performed using 8-16 week old adult male and female mice.

### Cell Lines

All hiPSC lines used in this study, both before and after gene editing, had a normal karyotype when analyzed by G-banding (Cell Line Genetics) and pluripotency was confirmed by staining for pluripotency markers as previously described.^87^ All hiPSC differentiations were performed under regulatory approval of the Institutional Review Board of Boston University. BU1 and BU3 control hiPSC lines were previously published.^28,29^ HPAH3 and HPAH4 hiPSCs lines were generated for the present study by reprogramming dermal fibroblasts from 2 donors with hereditary pulmonary arterial hypertension. Dermal fibroblasts from these donors were originally cryopreserved and archived at Vanderbilt University as part of a large, previously published, kindred of individuals with *BMPR2*-related pulmonary hypertension^7^. All protocols for rhe procurement of the HPAH3 and HPAH4 samples was approved by the Institutional Review Board of Vanderbilt University. Reprogramming methods are detailed below. Human umbilical vein endothelial cells (HUVECs) were obtained from Lonza (#C2519A). Note that all cell culture was performed in the presence of Primocin (InvivoGen, #ant-pm-2) and cultures underwent routine mycoplasma testing.

## Method Details

### Reprogramming to generate HPAH disease-specific human induced pluripotent stem cells (hiPSCs)

Reprogramming of dermal fibroblasts to generate HPAH3 and HPAH4 hiPSC lines employed protocols we have previously described, employing an excisable (floxed) single lentiviral “stem cell cassette” (hSTEMCCA) that encodes the 4 human reprogramming factors *OCT4*, *KLF4, SOX2*, and *cMYC*.^87^ Candidate hiPSC clones were screened by Southern blot to identify those carrying a single integrated copy of hSTEMCCA and then characterized by colony morphology and immunostaining against stage-specific embryonic antigen 4 (SSEA-4), TRA1-70, and TRA1-81 (Sigma, ES Cell Characterization Kit, #SCR001). The hSTEMCCA vector was then excised as previously described^87^ through transient transfection of a vector to express both Cre and puromycin resistance (pHAGE2-EF1a-Cre-IRES-Puro-W plasmid) using the TransIT-HeLaMONSTER^TM^ Transfection Kit (Mirus, MIR2904). Cells underwent puromycin selection using 1ug/mL for 2 days and then the antibiotic was removed and colonies were expanded. Single hiPSC clones were picked and screened by PCR for hSTEMCCA excision. hiPSC lines were accommodated to 2D Matrigel (Corning, #CB356278) and grown in mTesr1 medium (STEMCELL Technologies, #85850). hiPSCs were passaged by treatment with Gentle Cell Dissociation Reagent (STEMCELL Technologies, #100-0485). for 3 minutes and manually dissociated by scraping and gentle pipetting then transferred to a new 2D-coated Matrigel plate at a dilution of 1:10.

### CAS9-mediated gene editing to correct the *BMPR2* pathogenic variant

The HPAH3 and HPAH4 hiPSC lines harbored the heterozygous pathogenic *BMPR2* c.354 T>G, p.(C118W) variant in the third exon and underwent gene correction using CAS9-mediated gene editing. This was performed using the pSpCas9(BB) vector (Addgene #62988) to co-express Cas9 protein and the following guide RNA (gRNA): 5’-AAGGTTGAAGCCAGGCGTGA-3’ that targets a proto-spacer adjacent motif (PAM) within intron 3. The donor vector for homology directed repair (HDR) contained an 830 bp 5’ homology arm that included the corrected *BMPR2* exon 3 sequence and extended into the third intron. The 5’ homology arm was then followed by a puromycin resistance cassette flanked by loxP sites to be inserted in the third intron and then an 831 bp 3’ homology arm. To introduce the pSpCas9(BB) vector and HDR plasmid by nucleofection, hiPSCs were pre-treated with 10uM Y-27632 (Fisher, 12-545-0) for 3 hours, dissociated to single cells using Accutase (Sigma, #A6964) for 5-7 minutes to attain a single cell suspension, passed through a 30um strainer to remove clumps, and then pelleted at 200g for 4 minutes. Cells were then resuspended in 100ul of P3 Solution+Supplement (Lonza #V4XP-3012) with 2ug of pSpCas9(BB) and 3ug of HDR plasmid and underwent nucleofection in the Lonza 4D Nucleofector X system (AAF-1003x) using program CB-150. Nucleofected cells were cultured in media with Y-27632 for 24 hours followed by normal media conditions to allow colony expansion. Once colonies were visible (∼1 week), hiPSCs were then challenged with puromycin at 1ug/mL and surviving colonies were chosen for expansion. To excise the puromycin resistance cassette, resistant lines were then transfected with the pHAGE2-EF1a-Cre-IRES-Neo-W plasmid using the TransIT-HeLaMONSTER^TM^ Transfection Kit and underwent neomycin selection for 2 days. Surviving colonies were expanded and screened by PCR, the appropriate band was extracted using the QIAquick Gel Extraction Kit (Qiagen, #28704), and the sample was sequenced via Sanger sequencing to confirm successful gene correction and puromycin cassette excision. The following primers were used to amplify the target genomic region:

BMPR2_For: 5’-TTCCACCTCCTGACACAACA-3’
BMPR2_Rev: 5’-CGCTTTCGGATTTCAGGTGT-3’

### Directed differentiation and expansion of hiEndos

Directed differentiation of hiPSCs to generate hiEndos was performed through modification of previously published protocols,^26,30^ with modifications detailed in the results text and further detailed below. First, hiPSCs were pre-treated in 10uM Y-27632 for 1 hour and incubated in Gentle Cell Dissociation Reagent at 37C for 10-15 minutes until individual cells appeared rounded, but not yet dissociated from the plate. The Gentle Cell Dissociation Reagent was removed and cells were manually lifted in 1 mL of mTesr1 with 10uM Y-27632/well (6-well plate) using a cell scraper and gently pipetted to generate a single-cell suspension. 1×10^6^ cells were plated per well of a 6-well plate in 2mL of mTesr1 with 10uM Y-27632. After 24 hours, the medium was replaced with 1mL/well of Phase 1 medium: STEMdiff^TM^ APEL^TM^ 2 Medium (STEMCELL Technologies, #05275), 25ng/mL rhBMP4 (R&D Systems, #314-BP), 50ng/mL rhFGF2 (R&D Systems, #233-FB), and 6uM CHIR99021 (Tocris, #4423). After 24 hours in phase 1 medium, the medium was replaced with 2mL/well of Phase 2 medium: STEMdiff^TM^ APEL^TM^ 2 Medium, 25ng/mL rhBMP4, and 50ng/mL rhFGF2. After 24 hours, the medium was replaced with 2 mL/well Phase 3 medium: STEMdiff^TM^ APEL^TM^ 2 Medium, 200ng/mL rhVEGF-165 (R&D Systems, #293-VE), and 10um DAPT (Tocris, #2634/50). Cells were incubated in Phase 3 medium for 72 hours with media changes every 24 hours. On the final day of differentiation, cells were washed with 1x PBS and incubated in 1mL of Gentle Cell Dissociation Reagent for 5 minutes at 37C followed by 0.5 mL of 0.25% trypsin (Gibco, #25200056) for 3-5 minutes at 37C to generate a single cell suspension. The trypsin was inactivated in 0.5 mL of fetal bovine serum (FBS, Gibco, #16141079) and cells were pelleted at 300xg for 5 minutes to prepare for magnetic activated cell separation (MACs) to isolate endothelial cells.

The cell pellet was then incubated with anti-human CD31 (1:10, Miltenyi, #130-091-935) and anti-human CD144 microbeads (1:10, Miltenyi, #130-097-857) diluted in EGM2 medium (50uL total volume per well of a 6-well plate, Lonza, #CC-3162) for 20 minutes. The cells were then washed in 5ml of EGM2 medium and pelleted at 300xg for 5 minutes, resuspended in 1mL of EGM2 medium, and loaded on a pre-washed (3mL of EGM2 medium) LS MACs column (Miltenyi, #130-042-401) that was set in the MidiMACs^TM^ Separator (Miltenyi, #130-042-302) affixed to the MACS MultiStand (Miltenyi, #130-042-303). The MACs column was then washed three times with 3mL of EGM2 medium, removed from the MidiMACs Separator, and eluted into a fresh tube with 8mL of EGM2 medium to isolate the CD31/CD144 double positive hiEndos. Cells were then plated on 2D Matrigel-coated wells in medium composed of a 1:1 mixture of Phase 3 medium and ‘SD’ medium (EGM2 supplemented with 10uM DAPT and 10uM SB431542; Tocris, #1614; 2mL/well of a 6-well plate). The medium was switched to 100% SD medium the next day for cell maintenance and expansion with medium being changed every 24 hours. To passage hiEndos, cells were washed with 1x PBS, treated with Gentle Cell Dissociation Reagent at 37C for 5 minutes, and then 0.25% trypsin for 3 minutes at 37C to generate a single cell suspension. The trypsin was inactivated with equal parts FBS, pelleted at 300xg, and then split at a ratio of 1:3-1:4 into a new 2D Matrigel-coated plate. hiEndos may intermittently require re-purification via MACs in the first few passages should fibroblasts become visible in the culture.

### HUVEC Culture

Human umbilical vein endothelial cells (HUVECs) were cultured in EGM2 medium with daily medium changes. To passage, HUVECs were washed with 1x PBS and incubated with 0.25% Trypsin for 3-5 minutes at 37C to generate a single cell suspension. The trypsin was deactivated with 1 volume of FBS and the cells were pelleted at 300xg for 5 minutes. Cells were re-plated at a density of 1:3-1:6.

### Capillary Tube Formation Assay

Endothelial cells were plated at a density of 40,000 cells per well of a 48-well plate with wells prepared with 100ul of solidified 3D Matrigel (Corning, #356231). Cells were plated in 500ul of EGM2 medium in triplicate and imaged 12-18 hours after plating via phase contrast microscopy.

### Fluorescence activated cell sorting (FACs) and flow cytometric analysis

Single-cell suspensions of cells were prepared as described above for either cells in vitro or isolated from mouse lungs. Cells were pelleted to 300xg for 5 minutes and incubated with the indicated antibodies diluted in 1x PBS with 2% FBS (PBS+) at a total volume of 100ul for 30 minutes on ice. The cells were then washed with 1mL of PBS+ and centrifuged at 300xg for 5 minutes. The cells were then resuspended in 300uL of PBS+ with the viability exclusion dye DRAQ7 (1:500, abcam, #ab109202) and analyzed on a Stratedigm S100EXi or sorted using Beckman Coulter MoFlo Astrios. The following antibodies were utilized for flow cytometry and FACS: BV421-conjugated mouse anti-human CD31 (1:100, BD Biosciences, #564089); PE-conjugated mouse anti-human CD144 (1:20, BD Biosciences, #560410), BV421-conjugated mouse IgG1 k isotype control (1:100, BD Biosciences, #562438), PE-conjugated mouse IgG1 k isotype control (1:20, BD Biosciences, #555749).

### Acetylated low-density lipoprotein uptake assay

hiEndos were incubated in SBD medium supplemented with 5ug/mL of Alexa 594 conjugated acetylated low-density lipoprotein (594-AcLDL, ThermoFisher, #L35353) for 4 hours at 37C. The cells were then washed with 1x PBS and imaged by phase contrast and fluorescent microscopy using the Keyence BZ-X710 Cells were then dissociated by first washing with gentle cell dissociation reagent for 5 minutes at 37C followed by incubation with 0.25% trypsin for 3 minutes at 37C to generate a single cell suspension. The cells were then stained with a BV421-conjugated mouse anti-human CD31 antibody and analyzed on a Stratedigm S100EXi as above.

### hiEndo Growth Assays

For each condition, hiEndos were plated in duplicate wells of a 6-well plate at a starting cell number of 150,000 hiEndos and grown in the presence of vehicle, 10uM SB431542, 10uM DAPT, or combined SB431542+DAPT (each 10uM) in EGM2 medium. The medium was changed daily. When confluent, cells were dissociated as described above, a total cell count was obtained, and a subset of cells were stained with the BV421-conjugated mouse anti-human CD31 and PE-conjugated mouse anti-human CD144 antibodies and analyzed by flow cytometry on the Stratedigm S100EXi. hiEndos were replated at a concentration of 1:3 at each passage. To obtain the total hiEndo yield at each passage, the total cell number was multiplied by the proportion of CD31/CD144 double positive cells as determined by flow cytometry. hiEndos were grown until either there were no more CD31/CD144 double positive cells remaining in the culture, the cells underwent growth arrest, or passage 10 was reached at which point the assay was stopped.

### Lentiviral transduction of hiEndos

hiEndos were transduced with either pHAGE-EF1aL-EGFP-W^44^ or pHAGE-EF1aL-dsRED-W^45^ lentiviruses at a multiplicity of infection (MOI) of 10. The cells were dissociated to a single cell suspension as described above and then replated in a 2D Matrigel-coated well with virus in SBD medium (half volume, 1mL for well in 6-well dish) supplemented with 5ug/mL polybrene (Tocris, #7711). The medium was changed to fresh SBD medium after 16-18 hours and cultured as described above. All vectors are available by request through www.kottonlab.com.

### RNA Extraction and reverse transcriptase quantitative PCR (RT-qPCR)

RNA was isolated using the RNAeasy Mini Kit (Qiagen, #74104) per manufacturer’s protocol. cDNA was generated using the TaqMan Reverse Transcription Reagents (ThermoFisher, #N8080234) using 100-1000ng of RNA per sample. qPCR was set up using the TaqMan Fast Advanced Master Mix for qPCR (Applied Biosystems, #4444558) per manufacturers protocol for a 12ul reaction and run on an Applied Biosystems Quantstudio 6 Flex using the fast protocol for 40 cycles. TaqMan probes used in these studies: ACTA2 (Hs00426835_g1), APLN (Hs00175572_m1), APLNR (Hs00270873_s1), CA4 (Hs00426343_m1), CD36 (Hs00354519_m1), CDH5 (Hs00901465_m1), COL1A1 (Hs00164004_m1), CXCR4 (Hs00976734_m1), DLL4 (Hs00184092_m1), EDNRB (Hs00240747_m1), EPHB4 (Hs01119113_m1), FCN3 (Hs00892390_m1), FOXA2 (Hs05036278_s1), FOXF1 (Hs00230962_m1), GPIHBP1 (Hs01564843_m1), HPGD (Hs00960590_m1), ID2 (Hs04187239_m1), KDR (Hs00911700_m1), KIT (Hs00174029_m1), KITLG (Hs00241497_m1), MYH11 (Hs00975796_m1), NR2F2 (Hs00819630_m1), PAX2 (Hs01057416_m1), PDGFRA (Hs00998018_m1), PECAM1 (Hs01065282_m1), SOX17 (Hs00751752_s1), T (Hs00610080_m1), TBX2 (Hs00911929_m1), TBX6 (Hs00365539_m1), TMEM100 (Hs00388033_m1). The VIC Eukaryotic 18S rRNA Endogenous Control (Applied Biosystems, #4319413E) was used as a loading control. Each sample was run in technical duplicates and experimental (different wells of the same differentiation) vs biological replicates (distinct differentiations) are indicated in each figure legend. Relative gene expression normalized to 18S signal was reported as fold changes relative to a comparator control condition specified in each figure, with fold changes calculated as 2^-ddCT with the indicated control as per Pfaffi et al.^88^ Samples that did not have a detectable CT value were assigned a value of 40 to allow for calculations.

### Cell Signaling Assays

For the BMP9 and VEGF signaling assays (Figure 2), hiEndos were plated at a concentration of 150,000 cells/well of a 12-well dish and treated with the indicate growth factors (or vehicle controls) in EGM2 medium for 7 days at the specified concentrations. The medium was changed daily. RNA was isolated as above and the expression of the indicated genes were analyzed by RT-qPCR. Of note, SB431542 was included in the medium for all assays unless indicated otherwise. For the BMP9 signaling assays involving the HPAH3 and HPAH4 lines (Figure 5), hiEndos were plated at a concentration of 250,000 cells/well of a 12-well dish in EGM2 medium without serum [EGM2(-)] supplemented with 10uM SB431542. After 16-24 hours, the medium was replaced with EGM2(-) with 10uM SB431542 and either 40ng/mL BMP9 or vehicle-only. RNA was isolated after 24 hours of BMP9 vs vehicle-only treatment and gene expression was analyzed by RT-qPCR. For the phospho-SMAD experiments, hiEndos were plated at 150,000 cells per well of a 2D Matrigel-coated 8-well chamber slides (Millicell EZ Slide, Millipore Sigma, #PEZGS0816). The cells were grown to confluence then placed in EGM2(-) medium for 24 hours. The medium was then replaced with EGM2(-) with 40ng/mL of BMP9 or vehicle control and then fixed and stained 4 hours later as described below.

### hiEndo transplantation into the mouse lung

The hiEndos to be used for transplantation were lentivirally transduced to express either EGFP (pHAGE-EF1aL-EGFP-W^44^) or dsRED (pHAGE-EF1aL-dsRED-W^45^) as described above. The transduced hiEndos underwent 7-days of pre-treatment with EGM2 medium containing 40ng/mL BMP9 (BioLegend, #553104) and 10uM SB431542 unless otherwise indicated. The medium was changed daily and cells were passaged once during the pre-treatment phase. On day 4 of the pre-treatment phase, NSG mice aged 8-16 weeks were placed in the O2 Controlled InVivo Cabinet (Coy Labs) with oxygen levels set to 90% for 72 hours to induce hyperoxic injury. The cage was vented daily. On the day of transplantation, pre-treated hiEndos were dissociated to a single cell suspension as described above, washed with 1x PBS, and then resuspended in normal saline (150uL per 4,000,000 cell dose). The cells were kept under constant agitation on a rocker throughout the experiment to avoid cell clumping. Mice were anesthetized using isoflurane and hiEndos were injected intravenously via the retro-orbitall venous sinusy with 4,000,000 hiEndos in 150uL 1x PBS using a 27g tuberculin syringe (BD Biosciences, #14-829-9). Note that cells were drawn into the needle immediately before injection to avoid clumping in the syringe. Mice recovered on a warming pad before being transferred back to the mouse facility. Weights were measured daily for 72 hours post-transplantation during the recovery phase and dissected at the indicated times.

### Mouse lung single-cell dissociation and enrichment for hiEndos

Mice underwent euthanasia and perfusion through PBS injection to the right ventricle. The lungs were instilled with 1mL of 1xPBS via intratracheal injection three times to wash the tissue followed by instillation of 2mL of digestion buffer comprised of 9.5U/mL Elastase (Worthington Biochemical, #50-592-321), 20U/mL Collagenase type IV (ThermoFisher, #17104019), and 5U/mL Dispase (Gibco, #17105-041) in 1x PBS and the trachea was tied with a suture to contain the solution. Lungs were placed in ice cold 1x PBS and then each lobe was dissected out and placed in 3.5mL of digestion buffer and incubated at 37C on a rocker for 40 minutes. The tissue was then dissociated to a single cell suspension through sequential pipetting with a 10mL pipette, a 5mL pipette, and a P1000. The cell mixture was passed through a 70um then 40um filter to remove clumps and centrifuged at 300xg for 5 minutes. The pellet was incubated with 1mL of Red Cell Lysis Buffer (Sigma, #R7757) for 1 minute at room temperature, diluted in 5mL of 1x PBS, and pelleted at 300xg for 5 minutes. hiEndos were enriched from the cell mixture through MACS using anti-human CD31 and anti-human CD144 antibodies as described above. The resulting cells were stained with a B421-conjugated mouse anti-human CD31 antibody for FACS as described above.

### Competitive lung reconstitution assay

For the competitive lung reconstitution assays involving the comparison of pre-treatment strategies with the BU1 line (Figure 3), hiEndos derived from the same differentiation were transduced with either the pHAGE-EF1aL-EGFP-W^44^ or pHAGE-EF1aL-dsRED-W^45^ lentiviruses as described above. After growing to confluence, the hiEndos were dissociated to single cells, stained with the BV421-conjugated mouse anti-human CD31 antibody, and underwent FACS gating on human CD31 and either EGFP or dsRED to obtain a pure population of fluorophore-expressing hiEndos. The purified, fluorophore-expressing hiEndos then underwent differential pre-patterning strategies (i.e. BMP9+SB431542 vs SB431542 alone) for 7 days with an EGFP+ and dsRED+ population prepared for each pre-treatment strategy to enable a ‘color swap’ to control for fluorophore expression. Mice were placed in the hyperoxia chamber from days 4-7 of the pre-treatment phase as described above. On the day of transplantation, the hiEndos were processed and injected as described above with the following modifications. After generating a single cell suspension, the EGFP+ hiEndos from one condition were mixed 1:1 with the dsRED+ hiEndos of the other condition (and vice versa) to prepare enough cells to inject 4,000,000 hiEndos per animal (2,000,000 from each condition/dose). Before pelleting and resuspending in normal saline, the hiEndo mixture was analyzed on the Stratedigm S100EXi to ensure a 1:1 composition of EGFP to dsRED positive cells. The cell composition was adjusted as needed and reanalyzed until a 1:1 mixture was confirmed. The cells were then pelleted, resuspended in normal saline, and injected as above. After 2 weeks, the lungs were dissected, single cell dissociated, and hiEndos were enriched by MACS as described above. The resulting cells were stained with the BV421-conjugated mouse anti-human CD31 antibody and analyzed on the Stratedigm S100EXi to determine the proportion of EGFP+ to dsRED+ hiEndos (hCD31+) to obtain quantitative data related to the relative engraftment fitness of each cell population.

For the competitive lung reconstitution assays using the HPAH3 and HPAH4 lines, EGFP+ and dsRED+ hiEndos from each genotype were generated and purified as described above. All lines were patterned with BMP9 + SB431542 for 7-days as described above. On the day of injection, the EGFP+ *C118W/+* hiEndos were mixed 1:1 with dsRED+ *+/+* hiEndos (and vice versa) to prepare the resulting cell mixtures for injection.

### Immunofluorescence microscopy

For lung immunostaining, mice were euthanized and the lungs were inflated with 4% paraformaldehyde (PFA, Ted Pella, #18505) via intratracheal instillation under consistent 25 cmH_2_O pressure using a mounted syringe. The trachea was sutured shut prior to removing the intratracheal catheter and lungs were dissected out and placed in 4% PFA on a rocker at 4C overnight. The lungs were washed three times in 1x PBS on the 4C rocker for 30 minutes per wash and then cryoprotected in 30% sucrose in 1x PBS on the rocker at 4C overnight. The lobes were dissected and incubated in a 50% solution of OCT (Fisher, #23-730-571) and 30% sucrose in 1x PBS for 4-6 hours. The lung lobes were then embedded in a cryomold with OCT, frozen in isopropanol cooled with dry ice, and stored overnight at -80C. Tissue was then cryosectioned at a thickness of 30um, dried for at least 1 hour at room temperature, and washed 3 times in 1x PBS for 5 minutes per wash. The tissue was outlined with a PAP pen and blocking buffer [4% normal donkey serum (Jackson ImmunoResearch, #017000-121) in 1x PBS with 0.1% Triton-X 100 (Sigma-Aldrich, #X100)] was added for 1 hour at room temperature. The primary antibodies (see below) were diluted in blocking buffer and incubated on slides overnight at 4C in a hydration chamber. The slides were washed 3x in 1x PBS with 0.1% Triton-X 100 for 5 minutes per wash and then incubated for 1 hour in secondary antibodies (see below) diluted 1:500 in blocking buffer with Hoechst 33342 (1:500, ThermoFisher, #H3570). The slides were washed three times in 1x PBS with 0.1% Triton-X 100 for 5 minutes per wash and mounted with a coverslip using ProLong Diamond Antifade Mountant (Moleular Probes, #P36970).

For immunostaining of cultured cells, 150,000 hiEndos were plated in 2D Matrigel-coated 8-well chamber slides. When confluent, the cells were washed with 1x PBS and underwent fixation with 4% PFA for 10 minutes on ice. The wells were then washed three times with 1x PBS for 5 minutes each wash and then proceed with blocking, primary antibody, and secondary antibody incubations as described above.

The primary antibodies used in this study include: mouse anti-human CD31 (1:200, R&D Systems, #BBA7), rabbit anti-human CD144 (1:200, R&D Systems, #AF938), goat anti-human/mouse/rat CD31 (1:200, R&D Systems, #AF3628), hamster anti-mouse PDPN (1:500, ThermoFisher, #14-5381), rabbit anti-human phospho-SMAD1/5/9 (1:200, Cell Signaling, #13820), rabbit anti-human SMAD1/5/9 (1:200, abcam, #EPR25803-154). The secondary antibodies used in this study include (all at 1:500): Alexa Fluor 488 donkey anti-mouse IgG (ThermoFisher #A-21202), Alexa Fluor 546 donkey anti-rabbit IgG (ThermoFisher, #A10040), Alexa Fluor 647 donkey anti-mouse IgG (ThermoFisher, #A-31571), Alexa Fluor 647 donkey anti-goat IgG (ThermoFisher, #A-21447), and Alexa Fluor 546 goat anti-hamster IgG (ThermoFisher, #A-21111).

### Microscopy

Phase contrast microscopy was performed using the Keyence BZ-X710. Confocal imaging was performed using the Zeiss LSM 880 with Airyscan. Fluorescent microscopy was performed using the Nikon Eclipse Ni. Whole mount imaging of lungs was performed using the Nikon SMZ18 stereomicroscopy. ImageJ was used for fluorescent intensity quantification.

### *In vivo* intravascular labeling

To label lung-engrafted hiEndos *in vivo*, mice were retro-orbitally injected intravenously 2 hours before dissection with 100ug DyLight649-conjugated Ulex Europaeus Agglutinin I (UEAI-649, Vector Labs, #DL-1068-1) in 150uL total volume (diluted in 1x PBS). UEAI-649 binding was assessed by flow cytometry or immunofluorescence of cryosectioned tissue.

### RNA *in situ* hybridization

RNA *in situ* hybridization was performed using the RNAscope^TM^ Multiplex Fluorescent Reagent Kit v2 from Advanced Cell Diagnostics (ACD, #323100) using the 4-Plex Ancillary Kit (#323120) as per manufacturer’s protocol. The TSA Vivid Dyes (Tocris) 520 (#323271), 570 (#323272), and 650 (#323273) fluorescent reagents were used for visualization of the following RNAscope probes (ACD): Hs-PECAM1-no-XMm (#443471-C2), Hs-APLN (#44971-C1), Hs-APLNR (#450011-C3), Hs-VEGFA (#423161-C4), Hs-GJA4 (#856221-C1), EGFP (#400281-C2), DsRED (#481361-C3), and Hs-H19 (#400771-C1).

### Single cell RNA sequencing

Single cell suspensions were prepared as described above for cultured and transplanted hiEndos and stained with the BV421-conjugated mouse anti-human CD31 antibody. hiEndos were isolated for scRNA-sequencing via FACS on the Beckman Coulter MoFlo Astrios cell sorter for live hCD31+ cells as described above. The Chromium Single Cell 3’ System (10x Genomics) was used for scRNA sequencing via the Single Cell Sequencing Core at Boston University Medical Center according to the manufacturers protocol. Cell Ranger was used to demultiplex the results that were mapped using STARsolo to the human GRCh37/hg19 reference genome sequence that was extended to detect EGFP and dsRED transcripts. Raw gene expression matrices from 10x Genomics were processed using the Seurat v5.1.2^89^ package. Genes detected in fewer than five cells were excluded during object creation. Quality control metrics, including number of detected genes (nFeature_RNA), total UMI counts (nCount_RNA), and mitochondrial transcript percentage, were computed for each cell. Cells were filtered to remove low-quality profiles and potential doublets using thresholds of nFeature_RNA > 800, dataset-specific upper quantiles for nFeature_RNA and nCount_RNA, and mitochondrial content below 20%. Data were normalized using SCTransform with regression of mitochondrial percentage, followed by principal component analysis (PCA) on variable features. A shared nearest neighbor graph was constructed using the top 20 principal components, and unsupervised clustering was performed using the Louvain algorithm across multiple resolutions. Non-linear dimensionality reduction was performed using Uniform Manifold Approximation and Projection (UMAP) for visualization. Cell cycle phase (G1, S, G2/M) was assigned based on canonical gene signatures, and gene module scores were computed using Seurat’s implementation of the method described by Tirosh et al. 2016,^90^ incorporating various curated and published gene sets. Differential expression analysis between clusters was performed using the MAST^91^ framework, restricting testing to genes expressed in at least 25% of cells per cluster and applying a log fold-change threshold of 0.25; significant markers were defined using an adjusted p-value cutoff of 0.05. Quality control metrics, dimensionality reductions, and cluster annotations were visualized using Seurat, and heatmaps of marker genes were generated from scaled expression values. Gene set enrichment analysis was performed on cluster-specific differential expression results using pre-ranked gene lists based on average log fold-change. Genes were ranked per cluster, excluding non-finite values, and enrichment analysis was conducted using the fast gene set enrichment analysis (FGSEA)^92^ approach against multiple curated gene set collections including Hallmark Pathways and Gene Ontology Biological Processes (GOBP). Gene sets ranged in size from 5 to 500 genes, and enrichment scores were computed using a signal-to-noise–based ranking statistic. Enrichment results were computed independently for each cluster and gene set collection, and pathway-level significance was summarized per cluster for downstream interpretation and visualization. The scRNA-seq data discussed in this publication will be deposited in NCBI’s Gene Expression Omnibus at the time of final publication.

### Quantification and statistical analysis

Statistical details for each experiment are outlined in the corresponding figure legends. Unpaired and paired Student’s t-tests were used for data involving two groups and ANOVA was used when multiple groups were compared. The p value threshold to determine significance was set at p=0.05. Data for quantitative experiments are typically represented as the mean with error bars.

## Acknowledgements

We would like to extend our gratitude to the patients and their families who contributed samples for hiPSC reprogramming. We thank Dr. Jyh-Chang Jean and iPSC Core Manager, Dr. Marianne James, both of the Center for Regenerative Medicine of Boston University and Boston Medical Center, for assistance with reprogramming, gene editing, and biobanking. We thank Brian R. Tilton from the Boston University Flow Cytometry Core Facility for assistance with fluorescence activated cell sorting. We also thank Yuriy Alekseyev and Christopher Williams from the Boston University Single Cell Sequencing Core for assistance with library preparation and sequencing for our scRNA sequencing studies. Cartoons from the figures were generated using Biorender.com. This work was supported by NIH grants R01HL095993, P01HL170952, and U01HL148692 to DNK. hiPSC banking and resource sharing was supported by NHLBI grant NO175N92025D00035 to DNK. An NIH K08HL173561 and the Charles A. King Trust Post-Doctoral Fellowship FP01031866 to AMH, AMH was also supported by the T32GM007748.

## List of Supplemental Tables

**Table S1.** Gene sets used to analyze scRNA-seq datasets (in support of Figures 2, 4, 6, 7)

**Table S2.** Differential gene expression analysis by sample identity for hiEndo media conditions (in support of Figure 2)

**Table S3.** GSEA (GOBP) analysis of scRNA-seq data from hiEndo media conditions: EGM2 vs BVS and SD (in support of Figure S2)

**Table S4.** GSEA (GOBP) analysis of scRNA-seq data from hiEndo media conditions: BVS vs EGM2 and SD (in support of Figure S2)

**Table S5.** GSEA (GOBP) analysis of scRNA-seq data from hiEndo media conditions: SD vs EGM2 and BVS (in support of Figure S2)

**Table S6.** Differential gene expression analysis of hiEndos 5-weeks post-transplantation by clustering resolution 0.25 (in support of Figure 4)

**Table S7.** Differential gene expression analysis of HPAH4 scRNA-seq pre-transplant dataset by genotype (in support of Figure S7)

**Table S8.** Differential gene expression analysis of HPAH3 scRNA-seq pre-transplant dataset by genotype (in support of Figure S7)

**Table S9.** GSEA Hallmark analysis of HPAH4 scRNA-seq pre-transplant dataset by genotype (in support of Figure S7)

**Table S10.** GSEA Hallmark analysis of HPAH3 scRNA-seq pre-transplant dataset by genotype (in support of Figure S7)

**Table S11.** Differential gene expression analysis of HPAH4 post-transplant hiEndos by clustering resolution 0.5 (in support of Figure 7)

**Table S12.** Differential gene expression analysis of HPAH3 post-transplant hiEndos by clustering resolution 0.2 (in support of Figure 7)

**Table S13** Differential gene expression analysis of HPAH4 scRNA-seq post-transplant dataset by genotype (in support of Figure 7)

**Table S14** Differential gene expression analysis of HPAH3 scRNA-seq post-transplant dataset by genotype (in support of Figure 7)

